# A rare human centenarian variant of SIRT6 enhances genome stability and interaction with Lamin A

**DOI:** 10.1101/2021.12.13.472381

**Authors:** Matthew Simon, Jiping Yang, Jonathan Gigas, Eric J. Earley, Tory M. Schaff, Lei Zhang, Maria Zagorulya, Greg Tombline, Michael Gilbert, Samantha L. Yuen, Alexis Pope, Michael Van Meter, Stephan Emmrich, Jeehae Han, Seungjin Ryu, Archana Tare, Yizhou Zhu, Adam Hudgins, Gil Atzmon, Nir Barzilai, Aaron Wolfe, Kelsey Moody, Benjamin A. Garcia, David D. Thomas, Paul D. Robbins, Jan Vijg, Andrei Seluanov, Yousin Suh, Vera Gorbunova

**Affiliations:** Department of Biology, University of Rochester; Rochester NY 14627, USA; Department of Obstetrics and Gynecology, Columbia University; New York, NY 10032, USA; Biostatistics and Epidemiology, RTI International, Durham, NC, USA; Department of Biochemistry, Molecular Biology and Biophysics and Institute on the Biology of Aging and Metabolism, University of Minnesota; Minneapolis, MN 55455; Department of Biochemistry and Biophysics, Perelman School of Medicine, University of Pennsylvania; Philadelphia, Pennsylvania 19104, USA; Department of Genetics, Albert Einstein College of Medicine; Bronx, NY, USA; Department of Biology, Faculty of Natural Sciences, University of Haifa; Haifa, Israel 31905; Ichor Therapeutics; 2521 US-11, Lafayette, NY 13084, USA

**Keywords:** Centenarians, Lamin, Longevity, SIRT6

## Abstract

Sirtuin 6 (SIRT6) is a deacylase and mono-ADP ribosyl transferase (mADPr) enzyme involved in multiple cellular pathways implicated in the regulation of aging and metabolism. Targeted sequencing identified a SIRT6 allele containing two linked substitutions (N308K/A313S) as enriched in Ashkenazi Jewish (AJ) centenarians as compared to AJ control individuals. Characterization of this SIRT6 (centSIRT6) allele demonstrated it to be a stronger suppressor of LINE1 retrotransposons, confer enhanced stimulation of DNA double strand break repair, and more robust cancer cell killing compared to the wild type. Surprisingly, centSIRT6 displayed weaker deacetylase activity, but stronger mADPr activity, over a range of NAD^+^ concentrations and substrates. Additionally, centSIRT6 displayed a stronger interaction with Lamin A/C (LMNA), which correlated with enhanced ribosylation of LMNA. Our results suggest that enhanced SIRT6 function contributes to human longevity by improving genome maintenance via increased mADPr activity and enhanced interaction with LMNA.

## INTRODUCTION

SIRT6 is a protein deacylase and mono-ADP-ribosylase (mADPr) enzyme that has diverse cellular functions, many of which are related to aging and longevity. SIRT6 knockout mice show premature aging and genomic instability (Mostoslavsky, Chua et al., 2006) while mice overexpressing SIRT6 display an extended lifespan (Kanfi, Naiman et al., 2012, Roichman, Elhanati et al., 2021). Across mammalian species, SIRT6 is conserved and its activity has a strong positive correlation with maximum lifespan (Tian, Firsanov et al., 2019). At the molecular level, SIRT6 is involved in DNA repair, telomere maintenance, silencing of the repetitive elements including LINE1 retrotransposons, regulation of glucose homeostasis, inflammation and pluripotency (Etchegaray & Mostoslavsky, 2015, Kawahara, Michishita et al., 2009, Mao, Hine et al., 2011, Michishita, McCord et al., 2008, Mostoslavsky et al., 2006, Van Meter, Kashyap et al., 2014, Van Meter, Simon et al., 2016). In addition, SIRT6 is a noted tumor suppressor (Min, Ji et al., 2012, Sebastian, Zwaans et al., 2012, Van Meter, Mao et al., 2011).

Cells lacking SIRT6 are more susceptible to malignant transformation, while SIRT6 overexpression induces apoptosis in cancer cells, but not in normal cells (Van Meter et al., 2011).

SIRT6 is localized in the nucleus where it interacts with nuclear scaffold protein Lamin A/C (LMNA) (Ghosh, Liu et al., 2015), which is also implicated in human longevity. Mutations in LMNA cause a multitude of human genetic syndromes, many of them associated with premature aging, including Hutchinson Gilford progeria syndrome (HGPS) (De Sandre-Giovannoli, Bernard et al., 2003, Eriksson, Brown et al., 2003). Interaction with LMNA activates SIRT6 enzymatic activities (Ghosh et al., 2015). Early onset premature aging observed in SIRT6 deficient mice resembles HGPS, which led to the suggestion that abnormal SIRT6 localization and function in HGPS drives the disease pathogenesis (Ghosh et al., 2015). A coding change in *SIRT6,* SIRT6 D63H, leads to the loss of SIRT6 enzymatic activity and embryonic lethality (Ferrer, Alders et al., 2018).

The strong correlation between SIRT6 activity and longevity across species (Tian et al., 2019) and in genetically modified mice (Kanfi et al., 2012, Roichman et al., 2021) raises the question of whether higher SIRT6 activity may be associated with longer lifespan in humans. Non-coding genetic polymorphisms in the SIRT6 gene region were associated with human longevity in candidate SNP analyses (Hirvonen, Laivuori et al., 2017, Li, Qin et al., 2016, TenNapel, Lynch et al., 2014). To date no beneficial mutations in *SIRT6* associated with longevity have been functionally characterized.

In this study, we identified two novel variants of *SIRT6* enriched in a population of human centenarians. One of these variants, centSIRT6 possesses altered biological activities compared to the common allele. centSIRT6 allele demonstrates enhanced mono-ADP ribosylase activity, but a reduced deacetylase activity *in vitro*. This tradeoff in activity produces an allele that confers enhancement in DNA repair and enhanced suppression of transposable elements, as well as a resistance to oxidative stress. Additionally, the centenarian allele is more efficient at killing cancer cells. Functionally, we found that the centenarian allele shows an enhanced interaction with LMNA, which correlates with an increased ribosylation state of LMNA. Together, we identified a novel centenarian allele of SIRT6 that results in enhanced genome maintenance by shifting the balance between the deacetylation and mADPr activities of SIRT6.

## RESULTS

To examine whether variants of SIRT6 are associated with human longevity we performed targeted sequencing of SIRT6 locus in a population of 496 Ashkenazi Jewish (AJ) centenarians and 572 AJ controls (individuals without a family history of exceptional longevity). Table 1 shows all the SNPs identified. We observed association of rs350845 with living beyond 100 years (P = 0.049, **Table 1**). This SNP lies within a *SIRT6* intron and is an eQTL for *SIRT6* upregulation across 18 tissue types (**Fig EV1**). Interestingly, this SNP is in high linkage disequilibrium LD (r2 > 0.98) with two other eQTLs, rs350843 and rs350846, which also upregulate *SIRT6*.

In addition, two rare missense variants in perfect linkage were observed, rs183444295 (A313S) and rs201141490 (N308K), aka centSIRT6, which had nearly double the allele frequency among centenarians (1.0%) compared to both study controls (0.55%) as well as a separate AJ cohort within GnomAD which did not contain centenarians (0.60%), although this difference was not statistically significant (P=0.7, P=0.5, respectively) due to lack of power.

Analysis of the entire GnomAD database (141,456 individuals of diverse ethnic backgrounds), uncovered an apparent enrichment of centSIRT6 allele pair among the 75+ age group compared to other age groups. Compared to other alleles at similar MAF (0.1 – 1%), the centSIRT6 allele was in the top 5^th^ percentile of 75+ enriched SNPs and in the 9^th^ percentile of all missense mutations (**Fig 1A, B**). We sought to characterize this missense mutation to discover its molecular impacts on SIRT6 function in the context of extreme longevity (**Fig 1C**).

**Figure 1:**
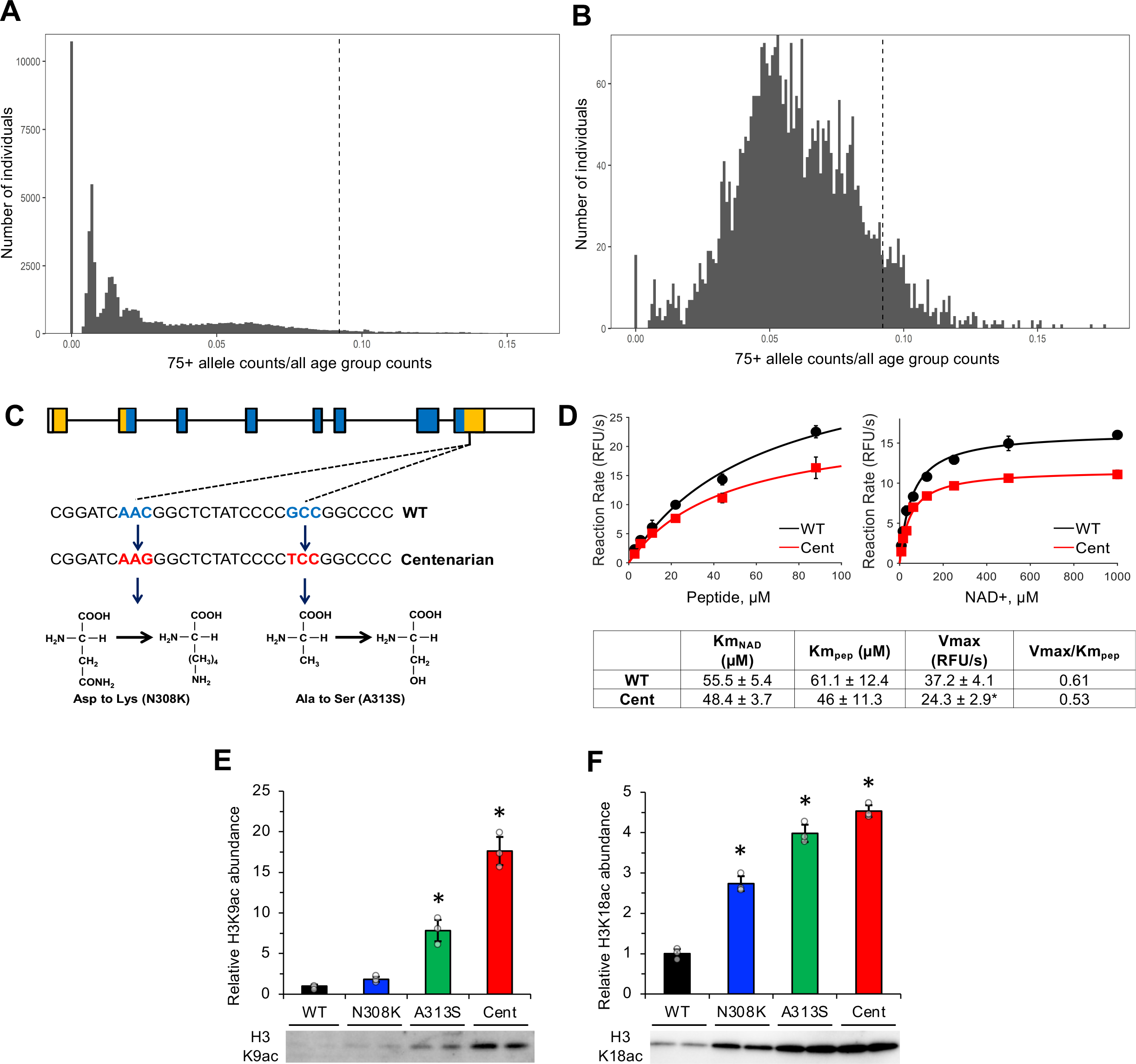
CentSIRT6 missense variant identified in AJ centenarians demonstrates lower deacetylation activity. **A, B** Histograms of normalized allele counts for SNPs in individuals 75+ years old compared to all age groups (N=125,748) across (**A**) entire chromosome 19, and (**B**) missense SNPs and indels only. Dotted vertical lines show centSIRT6. **C** The structure of the SIRT6 gene. centSIRT6 contains two missense mutations in the C-terminus N308K and A313S. White boxes represent UTRs, colored boxes represent protein coding regions, and blue boxes represent enzymatic domains. **D** Michaelis-Menten kinetic parameters as calculated by saturation curves using differential concentrations of myristoylated peptide or NAD^+^. Reactions were conducted in triplicate. **E, F** Deacetylase activity on H3K9 (**E**) and H3K18 (**F**) residues shows reduced activity in centSIRT6 allele. Designer histones saturated with the corresponding acetylated histone residue were incubated with purified SIRT6 and 1 mM NAD^+^ for 1 h prior to resolution by SDS-PAGE gel and staining with acetyl-specific histone antibodies. All reactions were conducted with an n=3; error bars show s.d. Statistics were calculated using Students *t*-test, two tailed. Asterisk indicate *p*<0.05.

### centSIRT6 demonstrates reduced deacetylase activity *in vitro*

SILAC analysis of HEK293 cells expressing different SIRT6 alleles showed no difference in the protein turnover rates between the WT and centSIRT6 (**Fig EV2A**). To assess the biochemical properties of the different SIRT6 alleles, we purified recombinant versions of each allele (**Fig EV2B**). Consistent with similar half-lives of wild type and centSIRT6 *in vivo*, we found no notable differences in the thermal stability of the SIRT6 alleles *in vitro*, suggesting that centSIRT6 does not grossly alter the folding or the ambient stability (**Fig EV2C**). As centSIRT6 mutations change the polarity and charge of the residues, we used SIRT6 FRET biosensors fused to the middle and the C-terminus of SIRT6 to assess for more subtle changes to protein structure that centSIRT6 mutations might cause (**Fig EV2D-G**). The centSIRT6 decreased FRET by ΔE = 6.4 ± 0.1 % relative to the wild type SIRT6, corresponding to an increase in the donor-acceptor distance R by 3.0 ± 0.4 Å. This increase indicates centSIRT6 mutations shift the SIRT6 structure toward a more open conformation.

We then tested SIRT6 deacylase activity (Jiang, Khan et al., 2013) on a myristoylated peptide to determine K_m_ for NAD^+^ and peptide for wild type and centSIRT6. While the centenarian allele displayed a slight reduction in activity (V_max_) compared to the wild type (*p* = 0.02) (**Fig 1C**), K_m_ was not significantly different between wild type and centSIRT6 for both substrates. Quenching of SIRT6 intrinsic Trp fluorescence by NAD^+^ binding also showed that centSIRT6 has similar affinity for NAD^+^ to the wild type allele (**Fig EV2H**). Overall, these data indicate that centSIRT6 confers a slight reduction in deacylase catalytic efficiency (as defined by V_max_ /K_m_) (**Fig 1C**).

The centSIRT6 displayed significantly lower deacetylase activity on recombinant nucleosomes with saturated H3K9ac and H3K18ac modifications (**Fig 1E, F**). Additionally, centSIRT6 allele displayed significantly slower histone deacetylase kinetics (**Fig 2A, B**). Similar to the synthetic histones, the centSIRT6 allele deacetylated both H3K9ac and H3K18ac residues on nucleosomes purified from HeLa cells, at a significantly reduced rate compared to the wild type SIRT6 (**Fig EV3A**). Taken together, the combined data from deacylase and histone deacetylase experiments indicate that the centSIRT6 variant has reduced deacylase/deacetylase activity.

**Figure 2:**
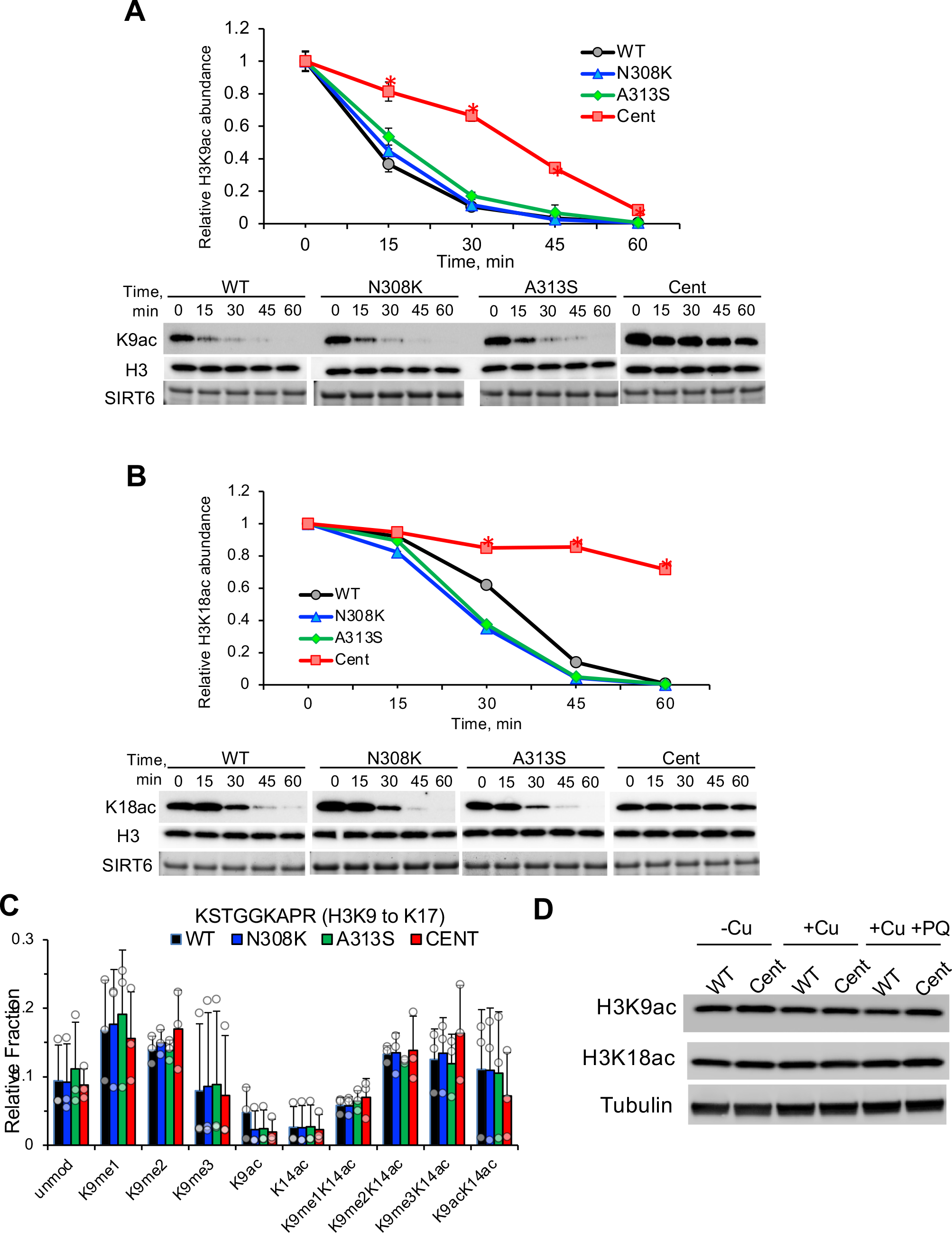
CentSIRT6 possesses reduced deacetylase activity. **A, B** Deacetylation kinetics of SIRT6 variants. Designer histones were incubated with purified SIRT6 and 5 mM NAD^+^, then resolved by SDS-PAGE and analyzed by immunoblotting with acetyl-specific histone antibodies. HeLa histone preps were also probed as shown in Fig EV3A. **C** Quantitative mass spec of histone H3 peptide purified from human cells expressing different SIRT6 alleles did not reveal a difference in acetylation levels. Relative fraction represents the portion of the peptide encompassing H3K9-17 compared to the summed total of all of the peptide quantitation values for the same region. The average and standard deviation of three different preps is plotted. Acetylation of other H3 peptides is shown in Supplementary Data 1. Histone preparation is shown in Fig EV3D. **D** Whole cell histone H3 acetylation levels in cumate-inducible SIRT6 human fibroblasts, assessed by Western blot. Cu, cumate; PQ, paraquat. Cumate dosage required for equivalent SIRT6 protein abundance was determined by Western blot, and administered accordingly to respective cell lines (Fig EV3B, EV3C).

To test the effect of the centSIRT6 allele on histone deacetylation activity *in vivo*, we generated telomerase-immortalized human HCA2 fibroblast cell lines in a SIRT6 KO background containing a cumate switch promoter driving expression of SIRT6 WT, N308K, A313S, and the centSIRT6 alleles (cumate SIRT6 fibroblasts). Each cell line was induced with cumate to drive equal levels of SIRT6 expression (**Fig EV3B, EV3C**). We quantified histone post-translational modifications from purified histones (**Fig EV3D**) from these cells by mass spectrometry using peptide standards as described previously (Sidoli, Bhanu et al., 2016). Cells expressing the centSIRT6, N308K, and A313S SIRT6 alleles showed similar global histone post translational modification (PTM) levels at all sites known to be SIRT6 substrates including H3K9ac, H3K18ac, and H3K56ac (**Fig 2C; Data S1**). Similarly, global levels of H3K9ac and H3K18ac measured by Western blot did not show significant changes, even when challenged with paraquat-induced oxidative damage (**Fig 2D**). These results suggest that while centSIRT6 enzyme has reduced catalytic efficiency of deacetylation, its activity is sufficient to maintain normal steady state levels of PTMs *in vivo*.

### mADPr activity is enhanced in centSIRT6

In addition to its deacetylase activity, SIRT6 has mono-ADP ribosylation (mADPr) activity, able to mono-ADP ribosylate itself and other proteins. mADPr activity is critical to SIRT6 role in DNA damage response and LINE1 repression (Kaidi, Weinert et al., 2010, Mao et al., 2011, Van Meter et al., 2014). To assess the mADPr activity, two known SIRT6 substrates were used: PARP1 and SIRT6, itself. The centSIRT6 demonstrated an enhanced auto- ribosylation rate with biotin-labeled NAD^+^ than the wild type SIRT6 (**Fig 3A**). The single SNPs did not display a notable difference in self-mADPr compared to the wild type, suggesting the increase observed in the centenarian allele results from a synthetic interaction that is not cumulative between the two SNPs (**Fig 3A**). Following this, we utilized an antibody (Bonfiglio, Leidecker et al., 2020) with specificity to mADPr residues to examine the self-ribosylation efficiency using a titration of NAD^+^. centSIRT6 displayed an apparent NAD^+^ affinity (K_app NAD_) of 142±29 μM and maximal relative auto-ribosylation mADPr_max_ of 5.27 whereas wild type SIRT6 displayed K_app NAD_ =106±45 μM and mADPr_max_ of 2.48. Thus, the overall efficiency as defined by mADPr_max_/ K_app NAD_ was 0.037 for centSIRT6 compared to 0.023 for wild type SIRT6, which also equates to a nearly two-fold difference (**Fig 3B**). In *trans*, mADPr activity was assessed by incubation with human PARP1, which was previously shown to be ribosylated by SIRT6 (Mao et al., 2011). Similar to self-ribosylation, the centSIRT6 displayed higher PARP1 ribosylation activity (**Fig 3C**). While both single mutants had elevated activity, the A313S allele was more similar to the centSIRT6 than N308K (**Fig 3C**). Collectively, these data show that centSIRT6 displays enhanced mADPr activity.

**Figure 3:**
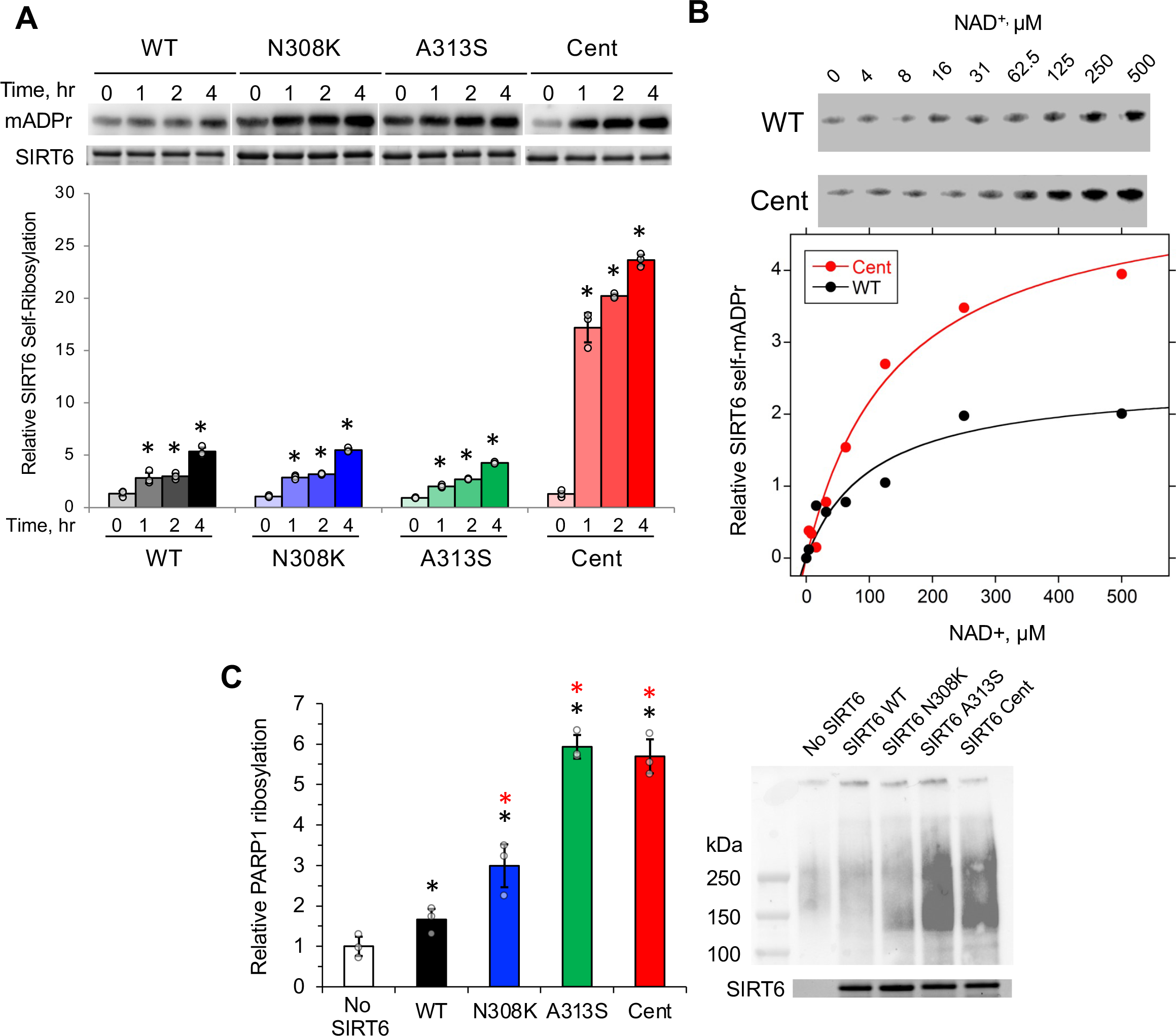
CentSIRT6 demonstrates enhanced mADPr activity. **A** Self-ribosylation of SIRT6 using biotin-labeled NAD^+^. Recombinant SIRT6 was incubated with NAD^+^ conjugated with a biotin residue and then run on an SDS-PAGE. Each allele was assessed relative to its 0hr time point and normalized to SIRT6 total protein loading controls. **B** Self-ribosylation of SIRT6 with titration of NAD^+^. mADPr-specific antibody(Bonfiglio et al., 2020) was used to detect ribosylated wild type SIRT6 and centSIRT6 proteins. Plot was fit the Michaelis-Menten equation with Kaleidagraph software. K_m NAD_ was 142±29 for centSIRT6 and 106±45 for wild type SIRT6 and maximal signal was ∼2x greater for centSIRT6 compared to the wild type SIRT6. **C** Activation of PARP1 by SIRT6 variants. SIRT6 protein was incubated with human PARP1 protein and then analyzed by immunoblotting with poly-ADPr antibody. Poly-ADPr activity of PARP1 results in a wide range of product size. Activity was assessed by quantifying poly-ADPr signal in whole lanes for each sample. All experiments were repeated at least three times; error bars show s.d. Significance was determined by Student’s *t*-test, two tailed. Asterisk indicate *p*<0.05. Black asterisks indicate significance over HPRT control. Red asterisks indicated significance over wild type SIRT6.

### centSIRT6 is more efficient at suppressing LINE1s and cancer cell killing

SIRT6 suppresses the expression of LINE1 retrotransposons via its mADPr activity (Van Meter et al., 2014). The loss of silencing of these elements contributes to age-related sterile inflammation and drives progeroid phenotypes in SIRT6 KO mice (Simon, Van Meter et al., 2019). To assess the capacity of the centSIRT6 allele to silence LINE1 transposons, we used the cumate-inducible fibroblasts expressing alleles of SIRT6. qRT-PCR analysis of both 5’- and 3’-biased regions of an evolutionarily active family of LINE1s both showed that the centSIRT6 allele enhances LINE1 suppression compared to the wild type SIRT6, as did the N308K allele (**Fig 4A**). The A313S allele showed a slight trend towards stronger suppression, but did not show a significant improvement over the wild type SIRT6 when assessed for the 3’ bias. These results show that centSIRT6 is more efficient at silencing LINE1 elements, which are functionally implicated in longevity.

**Figure 4:**
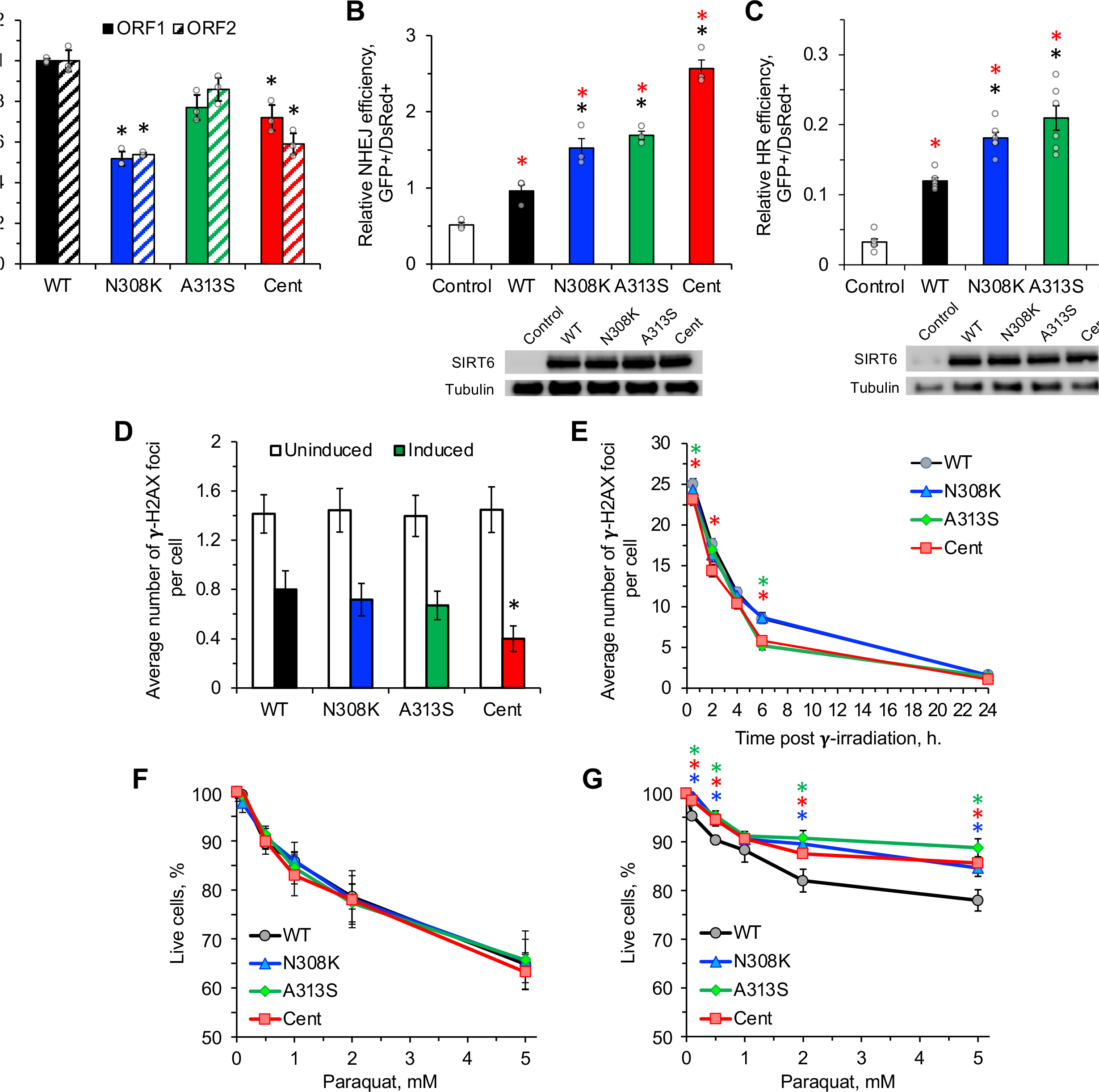
CentSIRT6 enhances LINE1 retrotransposon suppression and DNA repair. **A** qRT-PCR analysis of LINE1 expression in cumate-inducible SIRT6 fibroblasts. Primers assessed both 5’ (ORF1) and 3’ (ORF2) LINE1 sequences from the L1MdA1 family of active LINE1 retrotransposons. Assessment of both regions was conducted to mitigate contributions from partial insertion sequences in coding genes. **B, C** Stimulation of NHEJ and HR by SIRT6 variants. Reporter cell lines were co-transfected with SIRT6-expressing plasmid, I-Sce1 plasmid, and DsRed transfection control. After 72 hr recovery, reactivation of the GFP reporter was measured by flow cytometry. Stimulation of NHEJ or HR was calculated as radio of GFP^+^/DsRed^+^ positive cells. Representative FACS traces are shown in Fig EV5A. **D** Basal γH2AX foci in cumate-inducible SIRT6 fibroblasts. Foci were scored in at least 80 cells per condition. Representative images of the foci are shown in Fig EV4B. **E** DNA repair kinetics in cumate-inducible SIRT6 fibroblasts. Cells were grown on slides and irradiated with 2 Gy gamma radiation, followed by immunostaining for γH2AX. Irradiation was conducted when the cells were at 75% confluency on slides. Cells were fixed and foci scored at t=0.5 hr, 2 hr, 4 hr, 6 hr, and 24 hr post-irradiation. Foci were scored in at least 80 cells per genotype per timepoint. Representative images of the foci are shown in Fig EV5C. Asterisk indicates significant difference from the wild type SIRT6 (*p*<0.05). Color of the asterisk corresponds to the SIRT6 allele. **F, G** Oxidative stress resistance. Cumate-inducible SIRT6 fibroblasts were induced for SIRT6 expression and exposed to paraquat for 24 hours. Resistance was determined by apoptosis staining 48 hours after exposure. Representative FACS traces are shown Fig EV5D. All experiments were repeated at least three times. Error bars represent s.d. Significance was determined by Student’s *t*-test, two tailed, unless otherwise stated. Asterisk indicate *p*<0.05. Except in **E** and **G**, red asterisk indicates significant difference from control, and black asterisk indicates significant difference from the wild type SIRT6.

SIRT6 as a major regulator of DNA double strand break (DSB) (Mao et al., 2011). To quantify the differences between the SIRT6 alleles at promoting DSB repair, we employed *in vivo* GFP-based assays that measure nonhomologous end joining (NHEJ) and homologous recombination (HR) repair of a chromosomal DSB induced by I-SceI enzyme (Seluanov, Mao et al., 2010). Different alleles of SIRT6 were transiently expressed in the NHEJ and HR reporter telomerase-immortalized human foreskin fibroblast cell lines (Mao, Bozzella et al., 2008). We found that equivalent amounts of the centSIRT6 stimulated NHEJ 2.5-fold and HR 2-fold greater than the wild type SIRT6 (**Fig 4B, C; Fig EV4A**). Similarly, we observed in the cumate SIRT6 fibroblasts expressing centSIRT6 that basal levels of γH2AX foci were reduced by 50% relative to those observed in either single alleles or the wild type SIRT6 (**Fig 4D; Fig EV4B**). Taken together, these data indicate that the centSIRT6 allele elicits enhanced DNA DSB repair activity. We next compared whether different SIRT6 alleles have different effects upon stress. Using the cumate SIRT6 fibroblasts, we exposed the cells to γ-radiation and assessed the resolution of DSBs over time via γH2AX immunostaining. We observed that while both wild type and centSIRT6-expressing cell lines had similar levels of γH2AX foci immediately after exposure, the centSIRT6-expressing cells showed an improved recovery rate (**Fig 4E, Fig EV4C**). Furthermore, we found that the centSIRT6-expressing cells were more resistant to oxidative stress-induced apoptosis (**Fig 4F, G, Fig EV4D**). In contrast, in the absence of cumate induction, all cell lines showed no significant differences in survival after oxidative stress (**Fig 4F**). Taken together, these results indicate that the centSIRT6 improves DSB repair and resistance to DNA damage.

To further confirm that centSIRT6 enhances genome maintenance we generated knock-in human embryonic stem cells carrying centSIRT6 allele. The two centenarian mutations were introduced using CRISPR CAS9, and confirmed by sequencing. We generated two independent knock-in clones and differentiated them into MSCs. The MSCs were then transfected with linearized NHEJ and HR GFP reporters (Mao et al., 2008). Cells harboring centSIRT6 allele showed higher efficiency of repair (**Fig EV5A, EV5B**). These cells were also showed increased survival and lower number of 53BP1 foci upon treatment with methyl methanesulfonate (MMS), a potent DNA-damaging agent (**Fig EV54C, EV5D**). Interestingly, the centSIRT6 knock-in cells showed higher protein and mRNA levels of SIRT6 (**Fig EV5E, EV5F**), suggesting that centenarian mutations increase mRNA stability, in addition to modulating SIRT6 enzymatic activities. As it was impossible to compare equal levels of SIRT6 in CRISPR knock-in cells, we relied on cumate cells for the majority of our *in vivo* assays.

SIRT6 overexpression induces apoptosis in cancer cells but not in normal cells (Van Meter et al., 2011). In order to assess the tumor cell killing capacity of the centSIRT6 allele, we transfected SIRT6 alleles into two common tumor cell lines, HT1080 and HeLa cells. Expression of the centSIRT6 resulted in ∼2-fold fewer adherent surviving cells in both cancer cell lines compared to wild type SIRT6 (**Fig 5A**). Additionally, the centSIRT6 allele triggered at least 2- fold higher levels of apoptosis than the wild type SIRT6 allele in these cancer cell lines (**Fig 5B, C**). These data indicate that, centSIRT6 confers a more robust anti-tumor activity than the wild type SIRT6.

**Figure 5:**
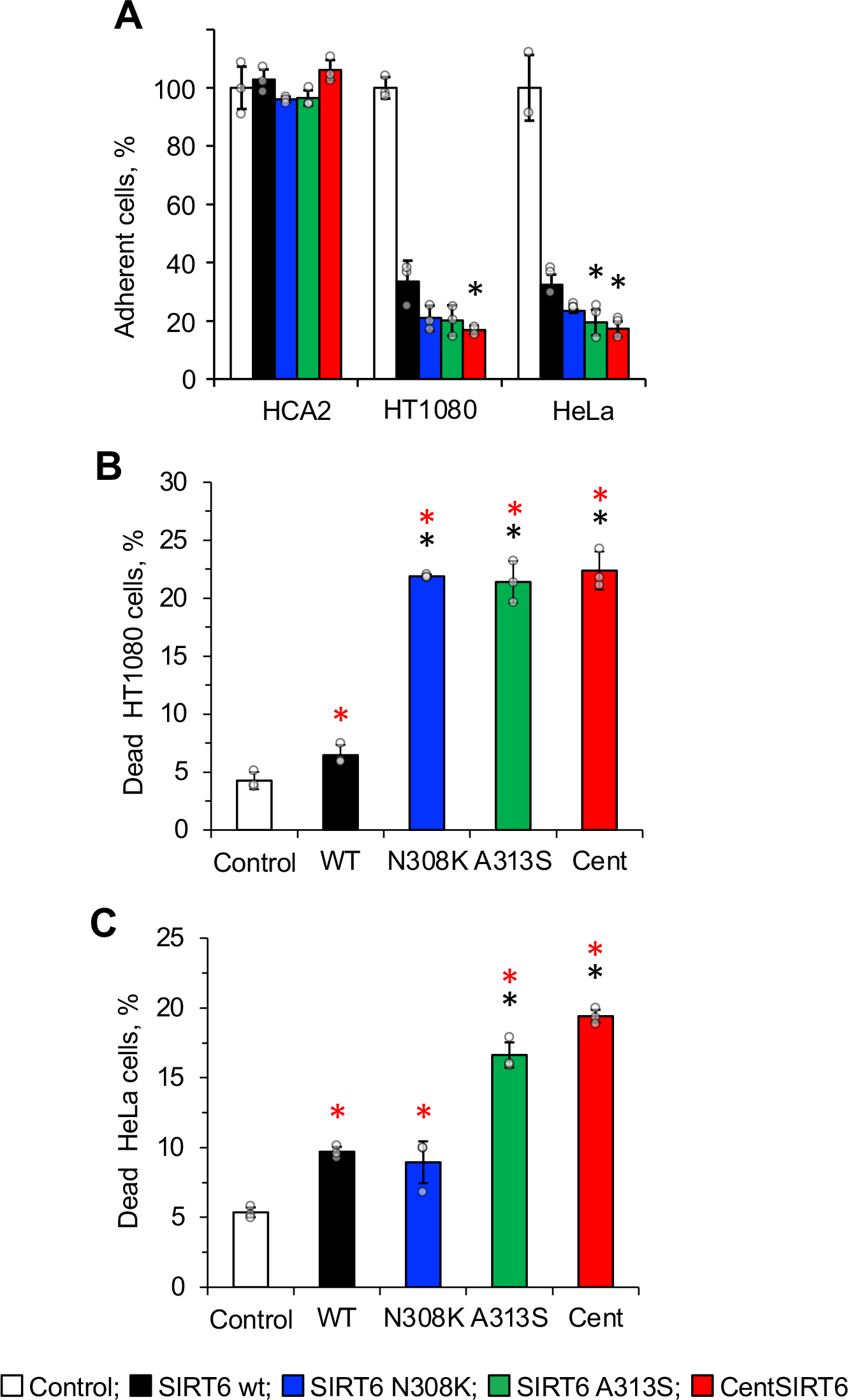
CentSIRT6 induces increase cell death in cancer cells. **A** Number of adherent cells after transfection with SIRT6 variants. Cells were transfected with SIRT6 plasmids encoding different SIRT6 alleles and cell numbers were counted after 72 hours. HCA2 are normal human foreskin fibroblasts. Asterisk indicates significant difference from the wild type SIRT6 (*p*<0.05). Color of the asterisk corresponds to the SIRT6 allele. **B**, **C** Apoptosis staining of cancer cell lines 48 hours after transfection. Cells were stained with Annexin V/PI and analyzed by flow cytometry. Significance was determined by two-way ANOVA. All experiments were repeated at least three times. Error bars represent s.d. Significance was determined by Student’s *t*-test, two tailed, unless otherwise stated. Asterisk indicate *p*<0.05. Red asterisk indicates significant difference from control, and black asterisk indicates significant difference from the wild type SIRT6.

### centSIRT6 displays higher affinity for LMNA

Since the centSIRT6 mutations are located in the flexible C-terminus of SIRT6 and may influence protein-protein interactions, we compared the interactomes of the centSIRT6 and the wild type alleles. We used antibodies against SIRT6 to immuno-precipitate SIRT6-interacting proteins from cumate SIRT6 fibroblasts and analyzed them by mass spectrometry with tandem-mass tags (TMT) to permit accurate quantitation. Several interacting proteins were enriched by the centSIRT6 allele (**Fig 6A; Data S2**). Most notable proteins showing stronger interaction with centSIRT6 than the wild type SIRT6 were LMNA and vimentin (VIME), which showed 38 and 40 peptides respectively. Both of these proteins are known to interact with SIRT6 (Ghosh et al., 2015, Gioutlakis, Klapa et al., 2017, Huttlin, Ting et al., 2015, Oughtred, Stark et al., 2019, Patil & Nakamura, 2005), and LMNA was shown to stimulate SIRT6 activity (Ghosh et al., 2015). No loss or gain of interactions was apparent in the centSIRT6 set, indicating that the effect of the novel allele could be in enhancing existing SIRT6 interactions.

**Figure 6:**
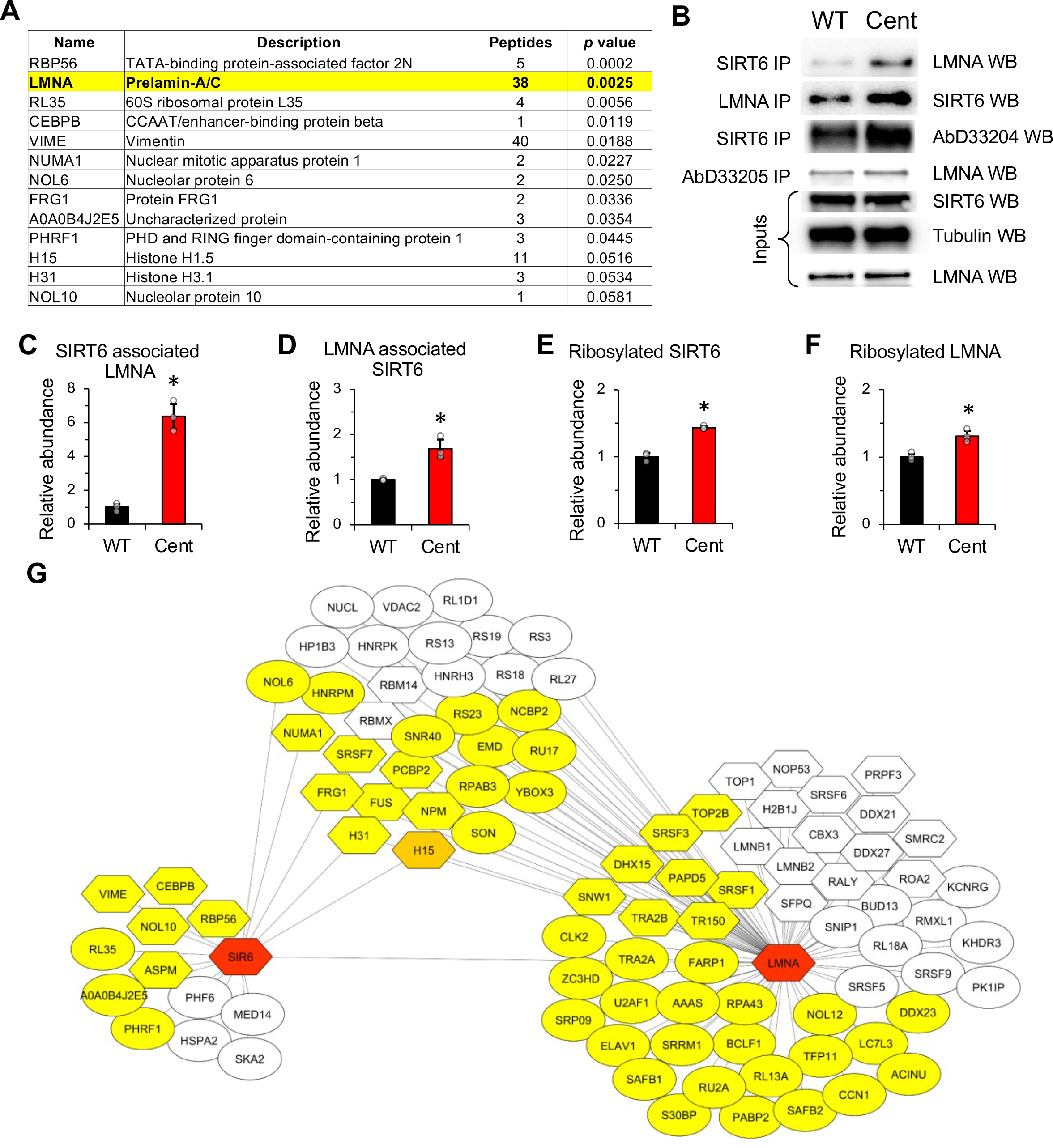
CentSIRT6 allele shows enhanced interaction with LMNA. **A** Proteins showing stronger interaction with centSIRT6 compared to the wild type SIRT6. Proteins were prepared from cumate-induced SIRT6 cell lines and immuno-precipitated with SIRT6 antibody; rabbit pre-immune serum was used as a control. Prior to analysis by mass spectrometry, samples were labeled with tandem-mass tags. **B** IP experiments on lysates from cumate-induced fibroblasts expressing wild type or centSIRT6 alleles with antibodies to SIRT6, LMNA, and mADPr. SIRT6 expression was induced 48 hours prior to IP. The IP experiments were repeated three times. One representative set of IPs is shown. **C** CentSIRT6 shows stronger interaction with LMNA compared to the wild type SIRT6. Quantification of the IP experiment shown in (**B**). SIRT6 IP followed by Western blot with antibodies to LMNA. **D** LMNA shows enhanced interaction with centSIRT6 compared to the wild type. Quantification of the IP experiment shown in (**B**). LMNA IP from cumate-inducible SIRT6 fibroblasts followed by Western blot with antibodies to SIRT6. **E** centSIRT6 shows enhanced mADPr. Quantification of the IP experiment shown in (**B**). SIRT6 IP from cumate-inducible SIRT6 fibroblasts followed by Western blot with antibody to mADPr residues(Bonfiglio et al., 2020) **F** LMNA shows enhanced mADPr signal in cells expressing centSIRT6. Quantification of the IP experiment shown in (**B**). IP with mADPr antibody(Bonfiglio et al., 2020) using extract from cumate-induced SIRT6 fibroblasts, followed by Western blot with antibodies to LMNA. **G** SIRT6 and LMNA are colored red and featured as central points in two opposing nodes of interactions. A third node (upper middle) shows interaction partners that are shared by SIRT6 and LMNA. Proteins whose interactions are enhanced by the centSIRT6 allele are colored yellow. Proteins that interacted equally with wild type and centSIRT6 alleles are uncolored. H1.5 is a special case and colored orange because it showed increased interaction with the centSIRT6 and decreased interaction with LMNA in the presence of the centSIRT6 allele. Proteins known to be ribosylated in previous reports are shown as hexagons. Error bars represent s.d. Significance was determined by Student’s *t*-test, two tailed. Asterisk indicate *p*<0.05

LMNA is a nuclear scaffold protein playing a central role in nuclear organization and is also implicated in aging. To confirm the mass spec data on SIRT6 and LMNA interactions, we performed co-IP with SIRT6 antibodies followed by Western blot with LMNA antibodies using cumate SIRT6 fibroblasts. We found that centSIRT6, indeed, associates more strongly with LMNA than the wild type SIRT6 allele (**Fig 6B, C**) and the effect was reciprocal as assessed by co-IP with LMNA antibodies followed by Western blot with SIRT6 antibodies (**Fig 6B, D**). Taken together these results demonstrate that centSIRT6 interacts more strongly with LMNA than the wild type SIRT6. Given the enhanced mADPr activity of the centSIRT6 allele, we next assessed the ribosylation status of LMNA using the mADPr antibodies. SIRT6 IPs performed on the cumate SIRT6 fibroblasts, using the mADPr specific antibody (Bonfiglio et al., 2020), showed that centSIRT6 was more ribosylated *in vivo* (**Fig 6B, E**); as demonstrated *in vitro* with the purified SIRT6 proteins (**Fig 3A**). IP with mADPr specific antibody (Bonfiglio et al., 2020) and subsequent Western blot with an anti-LMNA antibody revealed increased ribosylation of LMNA in the presence of the centSIRT6 allele (**Fig 6B, F**). LMNA has been reported to be ADP ribosylated *in vivo* on several residues (Adolph, 1987, Adolph & Song, 1985, Hendriks, Larsen et al., 2019).

We hypothesized that the centSIRT6 allele may influence the interaction of LMNA with other proteins. To test this, we immuno-precipitated LMNA from the cumate fibroblasts expressing either the wild type SIRT6 or centSIRT6 alleles and performed quantitative protein interaction analysis using TMT labeling followed by mass spectrometry. While many proteins showed similar interaction with LMNA, regardless of the SIRT6 allele present, a large group of proteins showed enhanced interaction with LMNA in the presence of centSIRT6 (**Fig 6G; Data S3**). LMNA-interaction partners enhanced by sentSIRT6 included ELAVL1, FUS, PCBP2, SAFB, SRSF1, SRSF3, TFIP11, THRAP3, BCLAF1, and U2AFBP all proteins that facilitate the coordination of the DNA damage response with RNA processing (Beli, Lukashchuk et al., 2012, Montecucco & Biamonti, 2013, Naro, Bielli et al., 2015, Vohhodina, Barros et al., 2017). Many of the proteins identified in SIRT6 and LMNA IPs are ADP-ribosylated (Bilan, Leutert et al., 2017, Hendriks et al., 2019, Jungmichel, Rosenthal et al., 2013, Leslie Pedrioli, Leutert et al., 2018, Martello, Leutert et al., 2016, Vivelo, Wat et al., 2017) (shown as hexagons in **Fig 6G**), suggesting that ADP-ribosylation by SIRT6 may play a role in mediating these interactions. Taken together, our data show that centSIRT6 is more efficient at promoting DNA repair and suppressing LINE1 elements, which is likely mediated by its enhanced mADPr activity and stronger interaction with LMNA.

## DISCUSSION

We demonstrated that a novel rare variant of SIRT6 discovered in a cohort of centenarians confers beneficial effects on SIRT6 protein function. centSIRT6 performed better at stimulating DNA repair, mitigating DNA damage, suppressing LINE1 transposons and killing cancer cells. Rare genetic variants are more likely to be deleterious than beneficial (Nelson, Wegmann et al., 2012), and are often associated with human disease. The only other coding SIRT6 human mutation, which had been functionally characterized, leads to embryonic lethality (Ferrer et al., 2018). SIRT6 activity is highly correlated with aging and it has been demonstrated that overexpression of SIRT6 results in increased lifespan in mice (Kanfi et al., 2012, Roichman et al., 2021). It has also been reported that SIRT6 from long-lived species can confer increased longevity in drosophila (Tian et al., 2019). Our findings suggest that increased SIRT6 activity also benefits human longevity.

Genetic association tests on human extreme longevity are uniquely challenging given the rarity of individuals and the lack of reliable statistical power to detect small effects. To date only two variants near APOE and FOXO3a have been associated with longevity in genome-wide scans (Broer, Buchman et al., 2015, Deelen, Beekman et al., 2011). For FOXO3a, this required a large-scale meta-analysis involving 6,036 longevity cases and 3,757 controls (Broer et al., 2015). Since there simply are insufficiently large numbers of centenarians in human populations, genetic association has limited utility in finding loci that promote healthy aging and longevity. In this study, we instead took a candidate SNP approach for *SIRT6* within a genetically isolated population of Ashkenazi Jews who exhibited extreme phenotype (>100 years old). The observed association for rs350845 which is linked to higher SIRT6 levels is nominally significant (P = 0.049). This association between higher SIRT6 expression and longevity is consistent with the data from Drosophila (Tian et al., 2019) and mouse (Kanfi et al., 2012) where SIRT6 overexpression resulted in lifespan extension. The missense variant centSIRT6 had twice the allele frequency in the AJ centenarians (1%) compared to controls (0.5%), however, we lacked power to detect statistically significant differences (P = 0.3). To further validate this finding, a follow up analysis of the GnomAD cohort of cohorts, which also contains AJ individuals, showed evidence of centSIRT6 enrichment among 75+ year-olds compared to all age groups. In addition to enrolling larger centenarian cohorts, future work may investigate epistasis and the co-occurrence of other SNPs in the genome with centSIRT6 and extreme longevity.

While increased SIRT6 activity correlates with longer lifespan in model organisms (Kanfi et al., 2012), (Tian et al., 2019), it has not been clear which of SIRT6’s enzymatic activities are central to longevity. The seemingly contradictory nature of the centenarian SIRT6 allele possessing decreased deacetylase activity came as an unexpected result. SIRT6 KO mice show several progeroid phenotypes and do not typically survive past 30 days (Mostoslavsky et al., 2006). In these mice, both the deacetylase activity and ribosylase activity are abrogated. In this study we have demonstrated that a centSIRT6 possesses a reduced deacetylase activity *in vitro* and showed non-discernable differences *in vivo*. The latter difference likely results from additional protein factors or PTMs present *in vivo* that help maintain SIRT6 deacetylation activity at close to the wild type levels, despite the weakened core enzyme. Interestingly, loss of SIRT6 deacetylation activity in humans or SIRT6 knockout in monkey lead to severe developmental phenotypes rather than premature aging, suggesting that, at least in primates, deacetylation activity of SIRT6 is required for development but not necessarily for adult longevity (Ferrer et al., 2018, Zhang, Wan et al., 2018).

The extent to which ADP-ribosylation functions of SIRT6 contribute to longevity remains largely unknown. Herein we report that the centSIRT6 allele displays significantly enhanced ribosylation activity against itself and PARP1 *in vitro* and also appeared enhanced *in vivo*. Further, we show that centSIRT6 confers improved aspects of SIRT6 function to living cells, including DNA DSB repair, LINE1 element suppression, and tumor cell suppression, all of which showed dependence on SIRT6 ADP-ribosylation functions previously. These results suggest that enhancement of ADP-ribosylation functions of SIRT6 contributes to increased longevity. While we cannot rule out that other aspects of SIRT6 activity, such as glucose metabolism and telomere maintenance, have been impacted by these mutations and may also play a role in the pro-longevity functions of this allele, those effects have been largely attributed to SIRT6 deacetylase activity which appears similar or reduced compared to the WT allele. It remains possible that SIRT6 ADP-ribosylation may play unanticipated roles in these processes as supported by recent reports of SIRT6 ADP-ribosylation of SMARCA2 and KDM2A (Rezazadeh, Yang et al., 2020, Rezazadeh, Yang et al., 2019). Consistent with this idea, SIRT6 mono-ADP ribosylation (and subsequent activation) of PARP1 plays a critical role in DNA DSB repair by facilitating recruitment of DNA repair proteins, such as MRN complex proteins (Haince, McDonald et al., 2008, Van Meter et al., 2016). Improvements in DSB repair efficiency, specifically those driven by SIRT6 activity, have been shown to correlate with longevity and healthspan (Roichman, Kanfi et al., 2017, Tian et al., 2019). Additionally, the ribosylation of KAP1 by SIRT6 and subsequent recruitment to transposable elements plays a critical role in silencing these opportunistic elements, which have been linked to age-related sterile inflammation and DNA damage (De Cecco, Ito et al., 2019, Simon et al., 2019, Van Meter et al., 2014). These same elements have been linked to inflammation driven by senescent cells, which are prevalent in aged systems (De Cecco et al., 2019). Further investigation of separation of function alleles of SIRT6 will be required to disentangle the contribution of its deacetylase/deacylase and ADP-ribosylation activities. Beyond its enhanced mADPr activity, centSIRT6 displays enhanced interaction with the nuclear scaffold protein LMNA. This result is especially significant for several reasons. LMNA functions as a key organizer of the nucleus, especially in maintaining heterochromatin at the nuclear periphery and LINE1 elements reside within lamina-associated domains (LADs) (Peric-Hupkes, Meuleman et al., 2010, Zullo, Demarco et al., 2012). Aberrant processing of LMNA results in human premature aging syndrome, Hutchison Gilford Progeria, while LMNA SNPs have been identified in human centenarians (Conneely, Capell et al., 2012, De Sandre-Giovannoli et al., 2003, Eriksson et al., 2003). Fibroblasts isolated from centenarians were also shown to have increased levels of pre- LMNA, suggesting that modulated functions of LMNA are associated with both premature aging and exceptional longevity (Ghosh et al., 2015, Lattanzi, Ortolani et al., 2014) found that the catalytic domain of SIRT6 interacts with LMNA. In addition, they found that many key aspects of SIRT6 activity, such as PARP1 ribosylation and activation, as well as chromatin localization in response to DNA damage were lost in LMNA^-/-^ MEFs. Our data suggest that a gain of function of SIRT6 may have the opposite effect of progerin on longevity. Remarkably, SNPs in LMNA were found in centenarians (Conneely et al., 2012). Fibroblasts isolated from centenarians were also shown to have increased levels of pre-LMNA, suggesting that modulated functions of LMNA are associated with both premature aging and exceptional longevity (Lattanzi et al., 2014).

In summary, we demonstrated that centSIRT6 displays reduced deacetylation activity *in vitro*, and has enhanced mADPr activity both *in vitro* and *in vivo*. This change in the balance of the two enzymatic activities leads to enhanced function of SIRT6 in DNA repair, LINE1 suppression and cancer cell killing, which are the activities that require mADPr activity of SIRT6. centSIRT6 also shows enhanced binding to LMNA which may further promote its function in DNA repair and chromatin organization via direct stimulation of SIRT6 enzymatic activity, and by coordinating interactions of LMNA with other components of LMNA complex (**Fig 7**). It would be interesting to model centSIRT6 in mice, to directly test whether this allele extends lifespan. However, the C-terminus of SIRT6 which harbors the centenarian mutations is not conserved in the mouse. SIRT6 and LMNA pathways are highly linked with the aging process, indicating that the SIRT6 ribosylation activity may be the more critical of the two enzymatic functions in regards to healthy aging and may aid LMNA to organize/control nuclear protein-protein and protein-RNA interactions. Given these results, there may be a benefit to enhancing SIRT6 activity, specifically the ribosylase activity. Molecules that impact SIRT6 activity, as well as its major co-enzyme NAD+, have been identified and hold potential as future methods of anti-aging interventions (Li, Bonkowski et al., 2017, Rahnasto-Rilla, Tyni et al., 2018, Rajman, Chwalek et al., 2018). Further refining the search for interventions that specifically target the mADPr activity of SIRT6 may yield more specific therapeutics to improve lifespan and healthspan.

**Figure 7:**
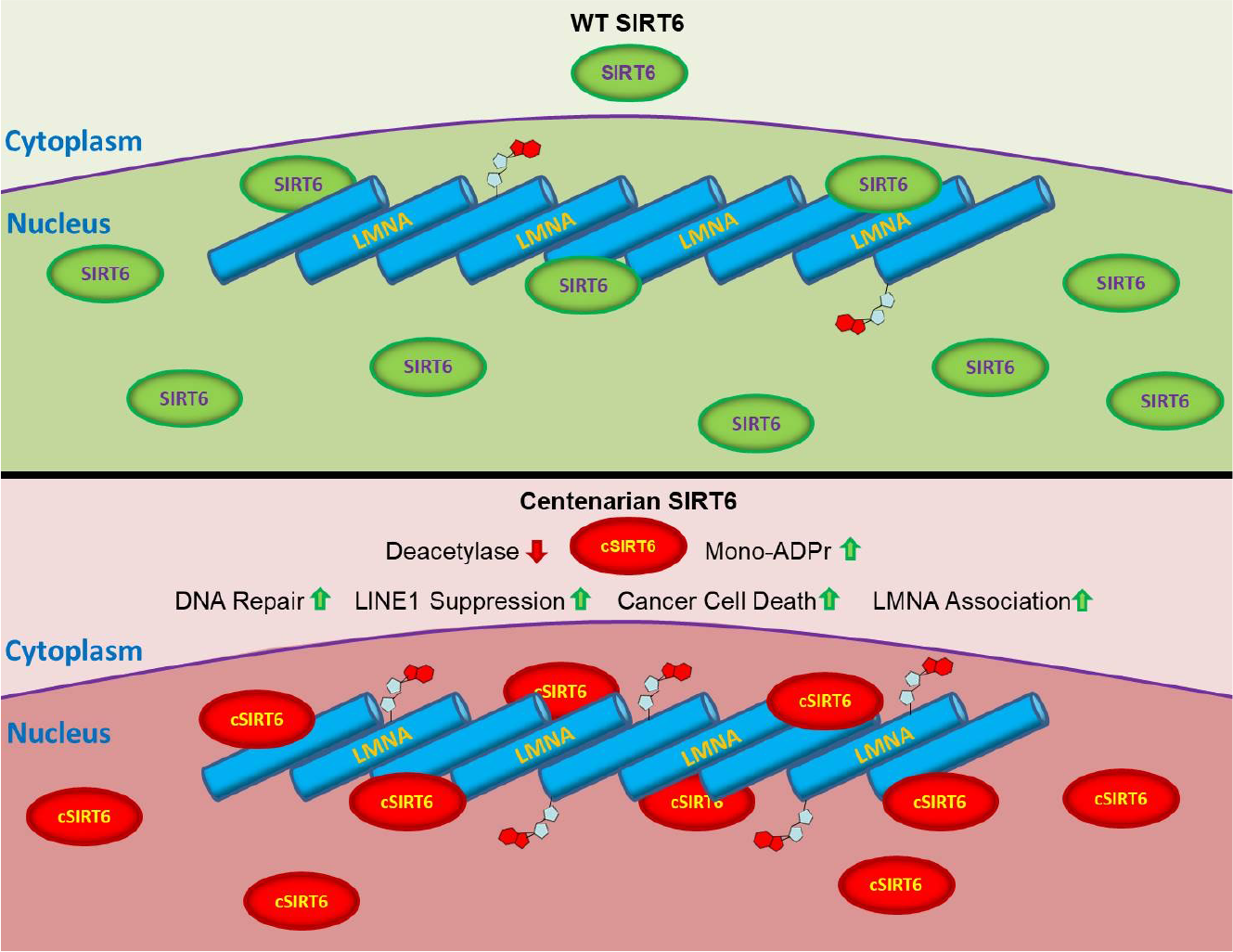
Altered molecular functions of centSIRT6. CentSIRT6 allele shows reduced deacetylation activity and enhanced mADPr activity. This results in enhanced DNA repair, improved LINE1 suppression and cancer cell killing. centSIRT6 shows stronger interaction with LMNA and enhances LMNA interactions with its partners.

## MATERIALS AND METHODS

### Human subjects

Ashkenazi Jewish population derived from an undetermined small number (estimated to be in the several thousands) of founders. External factors, including ecclesiastical edicts prohibiting all social contact with Jews, the Crusades, the establishment of the Pale of Settlement, numerous Pogroms, and ethnic bigotry, resulted in the social isolation and inbreeding of the Ashkenazi Jews. This history resulted in both cultural and genetic homogeneity and has made this population useful for identification of genetic traits. In our study, a centenarian is defined as a healthy individual living independently at 95 years of age or older and a control is defined as an individual without a family history of unusual longevity; parents of controls survived to the age of 85 years or less. The participants’ ages were defined by birth certificates or dates of birth as stated on passports. This study group consisted of 496 Ashkenazi Jewish centenarians and 572 Ashkenazi Jewish controls that were previously collected as part of longevity study at the Albert Einstein College of Medicine (Barzilai et al, JAMA, 2003).

Informed written consent was obtained in accordance with the policy of the committee on clinical investigations of the Albert Einstein College of Medicine, New York, NY. An additional 5,185 AJ controls were derived from GnomAD data base.

### Cell lines

HEK293 SIRT6 overexpression lines were generated by transfecting HEK293 cells with linearized CMV-SIRT6 plasmids via jetPRIME transfection reagent and selecting stably integrated clones. To generate normal human fibroblasts expressing WT and centenarian alleles of SIRT6 under control of cumate-inducible promoter (cumate SIRT6 fibroblasts), constructs containing SIRT6 alleles under control of cumate promoter were integrated in the genome of telomerase-immortalized SIRT6-KO human HCA2 foreskin fibroblasts via the PiggyBac Transposon Vector System. Endogenous SIRT6 was knocked out in these cells using CRISPR/Cas-9 (Tian et al., 2019). NHEJ and HR reporter assays were conducted using telomerase-immortalized HCA2 human fibroblast cell lines containing integrated reporter constructs (I9A and H15C) (Mao et al., 2008, Seluanov et al., 2010). HT1080 and HeLa cell lines were used to assess anti-tumor activity. Human H7 ESCs (WiCell Research) and their genetic modified derivatives were maintained on Matrigel in mTeSR Plus medium and used to generate MSCs.

### Cell culture

All cell lines were maintained in humidified incubators at 5% CO_2_, 5% O_2_, at 37°C. Cells were grown in Eagle’s minimum essential medium with 15% fetal bovine serum and 1x penicillin/streptomycin, with the exception of the HEK293, HT1080 and HeLa cells, which were culture in DMEM with D-Glucose and L-Glutamine. Human MSCs were cultured on Gelatin-coated plate in hMSC culture medium (αMEM (Gibco), 10% fetal bovine serum (GeminiBio), 1% penicillin/streptomycin (Gibco) and 1 ng/mL bFGF (ThermoFisher)). The cell lines are routinely tested for mycoplasma contamination.

### Sequencing of *SIRT6*gene in Ashkenazi Jewish Centenarians

The sequencing of SIRT6 gene was a part of the pooled target capture sequencing (capture-seq) project aiming to discover centenarian-enriched rare variants in genes involved in conserved longevity assurance pathways (Ryu, Han et al., 2018). SIRT6 was selected as a candidate longevity-associated gene. All SIRT6 variants discovered in the capture-seq of 1,068 AJ samples (496 centenarians vs. 572 controls) are provided in **Table I**.

**Table I.** List of all SIRT6 variants identified by capture sequencing.

| SNP ID | Alt | Ref | Variant Type | AA change | Current Study Centenarians |  |  | Current Study Controls |  |  |  | GnomAD Controls |  |  |  |
| --- | --- | --- | --- | --- | --- | --- | --- | --- | --- | --- | --- | --- | --- | --- | --- |
|  |  |  |  |  | Alt Count | Ref Count | MAF | Alt Count | Ref Count | MAF | P | Alt Count | Ref Count | MAF | P |
| rs350845 | G | A | intron | - | 749 | 151 | 0.17 | 962 | 138 | 0.126 | 0.009 | 8783 | 1367 | 0.135 | 0.007 |
| rs11554579 | C | T | synonym | R76R | 13 | 887 | 0.014 | 10 | 1090 | 0.009 | 0.296 | 91 | 9785 | 0.009 | 0.15 |
| **rs183444295** | **A** | **C** | **missense** | **A313S** | **9** | **891** | **0.010** | **6** | **1094** | **0.006** | **0.3** | **25** | **4281** | **0.006** | **0.267** |
| **rs201141490** | **C** | **G** | **missense** | **N308K** |  |  |  |  |  |  |  | **28** | **4826** | **0.006** | **0.268** |
| rs748302927 | T | G | intron | - | 1 | 899 | 0 | 0 | 1100 | 0 | 0.45 | 5 | 9375 | 5E-04 | 0.423 |
| rs992892719 | G | C | missense | R291P | 24 | 876 | 0.03 | 27 | 1073 | 0.025 | 0.777 | 0 | 6726 | 0 | <0.001 |
| rs200174862 | T | G | intron | - | 1 | 899 | 0 | 1 | 1099 | 9E-04 | 1 | 3 | 4777 | 6E-04 | 0.499 |
| rs200813796 | T | C | 3' UTR | - | 3 | 897 | 0 | 3 | 1097 | 0.003 | 1 | 11 | 4075 | 0.003 | 0.728 |

AJ, Ashkenazi Jewish; MAF, minor allele frequency; REF, reference allele; ALT, alternative allele; CON, controls; CENT, centenarians; P-values from Fisher exact test on allele counts.

Plots of *SIRT6* expression and eQTL status for rs350845 were downloaded from the GTEx website.

### GnomAD *centSIRT6* enrichment

GnomAD analysis was performed on the 2.1.1 release of combined exome and whole-genome sequences representing 125,748 exomes and 15,708 whole genomes. Chromosome 19 variants were downloaded from the GnomAD database, and since the centSIRT6 allele is rare (0.5% - 1%) we filtered for SNPs between 0.1% and 1% frequency. Our goal was to build a distribution of allele occupancy at the 75+ age group normalized against all age groups. While this is not a traditional association study per se, it does allow us to witness allele frequency bias in older vs. younger age brackets.

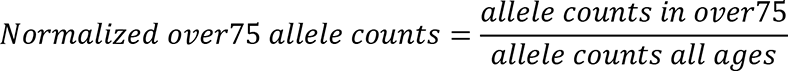

A normalized value of 0 represents a complete absence of the allele in 75+, and higher values represent greater occupancy of that allele in the 75+ age bracket compared to all ages. Alleles with a value of 1 were filtered from consideration, as these may have represented alleles only observed in cohorts comprised of older individuals. In total 66,441 SNPs were measured, of which 3,307 were missense.

### Gene editing at SIRT6 locus

CRISPR/Cas9-mediated knock-in was performed using Alt-R CRISPR-Cas9 System (IDT) and HDR Donor Oligos. Cas9 nuclease, sgRNA (GAATCTCCCACCCGGATCAA) targeting double variant site and single-strand DNA oligo donors (ssODNs) containing SIRT6 double variant were ordered from IDT. To generate SIRT6 double variant knock-in hESCs, 2×10^5 individualized hESCs were resuspended in 10 μL Resuspension Buffer R (Invitrogen) containing CRISPR ribonucleoproteins (Cas9 protein+sgRNA) and ssODNs and were then electroporated using NEON Transfection System (Invitrogen). After electroporation, cells were seeded on Matrigel-coated plates in mTeSR Plus with 1x RevitaCell Supplement (Gibco). After 48h expansion, cells were dissociated by Accutase and 10,000 cells were seeded on CytoSort™ Array (10,000 microwells, CELL Microsystem). Once cells were attached, microwells containing single colony were automatically picked and transferred to 96-well plate by CellRaft AIR System (CELL Microsystem). The expanded clones on 96-well plate were further genotyped by TaqMan genotyping assay (rs201141490, rs183444295, ThermoFisher) and Sanger sequencing.

### MSC differentiation

hESCs-derived embryoid bodies were first produced using AggreWell (Stem Cell technology) and were then plated on Matrigel-coated plates in hMSC differentiation medium (αMEM (Gibco), 10% fetal bovine serum (GeminiBio), 1% penicillin/streptomycin (Gibco), 10 ng/mL bFGF (ThermoFisher) and 5 ng/mL TGFβ (Thermofisher)) for around 10 days till fibroblast-like cells were confluent. These fibroblast-like cells were maintained in hMSC culture medium on Gelatin-coated 10cm dishes for two passages and were further sorted by FACS machine (BD FACS Aria II) to purify CD73/CD90/CD105 tri-positive hMSCs.

### Western blotting

Cells and reactions were collected using a 2x Laemmli solution and incubated on ice for 15 min, during which time the samples were passed through a large gauge needle several times and vortexed every 5 min. Samples were then spun at 14,000 RPM to remove debris and the supernatant transferred to a new tube. Samples were heated in boiling water for 20 min before being centrifuged at 14,000 RPM for 1min and loaded into a BioRad Criterion 4-20% gel. After transfer to PDVF membrane and blocked (5% dehydrated milk) for 2 hr at RT, membranes were incubated overnight antibodies in 2.5% blocking buffer at 4°C. Membranes were washed 3x with TBST for 10 min each before secondary antibody in 1x TBST was added for 2 hr incubation at RT. Membranes were washed 3x with TBST for 10 min and then imaged.

The following antibodies were used: H3 (Abcam ab500)-1:5000, H3K9ac (Abcam ab4441)- 1:1000, H3K18ac (Abcam ab1191)-1:1000, β-tubulin (Abcam ab6046)-1:10,000, SIRT6 (Cell Signaling #12486)-1:1000, Lamin A/C-1:1000 (Abcam ab108595, Millipore 05-714), γH2AX (Millipore 05-636), PARP1(Abcam ab227244)-1:1000, mADPr 1:500 (AbD33204 and AbD33205)(Bonfiglio et al., 2020).

### Immunoprecipitation

Cells were plated and grown with equivalent dosages of cumate for 48 hrs prior to IP. In brief, cells were collected via trypsin and centrifugation and then lysed on ice using IP Buffer (20 mM HEPES pH=8, 0.2 mM EDTA, 5% glycerol, 150 mM NaCl, 1% NP40) + COMplete protease inhibitor for 10 min. Lysate was sonicated at 25% output for 10 pulses and then debris pelleted by centrifugation at 4°C at 13000 RPM for 10 min. Supernatant was transferred to a new tube and precleared with A/G Sepharose beads at 4°C for 1 hr on a rotator. Beads were removed by centrifugation and transfer of cleared sample to new tubes. 50 μl of sample was reserved as input control. Samples were incubated overnight at 4°C with 3 μg of antibody (SIRT6-Cell Signaling) #12486; PARP1-Abcam ab227244; Lamin A/C-Millipore 05-714; AbD33204(Bonfiglio et al., 2020), AbD33205(Bonfiglio et al., 2020), then with 30 μl of 25% Sepharose for 2 hr at 4°C. Samples were spun down and beads washed 5x with IP buffer. Final resuspension with 100 μl IP buffer.

### DNA Repair Assays

Both NHEJ and HR efficiency were assessed as previously described(Seluanov et al., 2010). In brief, I-Sce1, SIRT6 or HPRT control, and dsRED plasmid were transfected into telomerase-immortalized human foreskin fibroblasts containing chromosomally integrated NHEJ or HR reporter cassettes (I9A or H15C) cells (Mao et al., 2008). Cells were allowed 3 days to recover prior to analysis by flow cytometry. Efficiency was calculated as the ratio of GFP events over dsRED events.

### Transfections

Transfections were carried out by plating cells at a density of 500,000 cells/10 cm plate two days prior to transfection. Transfections were carried out using the Amaxa Nucleofector with Normal Human Dermal Fibroblast transfection solution following manufacture’s protocol. For cancer cell transfections, jetPRIME transfection reagent was used to deliver plasmids into cells.

### Quantitative RT-PCR

Total RNA was isolated from cells at 80% confluence using Trizol Reagent and then treated with DNase. cDNA was synthesized using Superscript III (Life Technologies) cDNA kit with the OligodT primer. qRT-PCR was performed on the BioRad CFX Connect Real Time machine with SYBR Green Master Mix (BioRad) using 30 ng of cDNA per reaction with 3x reactions/sample. All primer sets were tested for specificity and efficiency. Actin was assayed using QuantumRNA Actin Universal primers (Thermo AM1720). LINE1 ORF1 Fwd- ATGGCGAAAGGCAAACGTAAG, Rev-ATTTTCGGTTGTGTTGGGGTG; LINE1 ORF2 Fwd GCAGGGGTTGCAATCCTAGTC, Rev- CTGGGTGCTCCTGTATTGGGT.

### Immunofluorescence and apoptosis

γH2AX and 53BP1 immunostaining was carried out as previously described(Mao et al., 2011). Anti- γH2AX (05-636) and anti-53BP1 (MAB3804) antibodies were purchased from Millipore. Apoptosis in fibroblasts was measured using the Annexin V Staining Kit (Roche).

### MMS treatment

3x10^4^ hMSCs were seeded on gelatin-coated 96-well plate and hMSCs were treated with DNA-damaging agents for 48 hours upon reaching 90% confluence. CellTiter 96 AQueous One Solution Cell Proliferation Kit (MTS assay, Promega) was used to measure cell viability according to the manufacturer’s protocol. DNA-damaging agents employed in this assay were as follows: Methyl methanesulfonate (0.125/0.25/0.5 mM, MMS).

### Gamma Irradiation

Cells were grown to 75% confluency prior to treatment on coated slides. A Cs-137 irradiator was used to deliver 2Gy of radiation to cells. Cells were transported in a 37°C container and media was replaced post exposure.

### Paraquat Treatment

Cells were maintained to 75% confluency, at which point fresh media lacking sodium pyruvate and containing paraquat was added. Cells were maintained for 24 hrs in treated media, followed by replacement with fresh media lacking sodium pyruvate. 48 hrs post treatment, the cells stained and evaluated for apoptosis.

### SIRT6 Protein Purification

His-tagged SIRT6 cDNA was cloned into pET11a vectors and transformed into Rosetta-Gami *E. coli* cells. Cells were grown in presence of antibiotics and then protein production was induced with 0.5mM IPTG 2 hrs before harvest. Cells were spun down and lysed on ice for 1h using a in a solution of 50 mM Tris-HCl (pH=7.5), 300 mM NaCl, 10% glycerol, and 10 mM imidazole with EDTA-free protease inhibitor (Sigma #P8849) and 1mg/ml egg white lysozyme followed by sonication with a Branson instrument. After removal of cell debris by centrifugation, lysate was incubated with Ni_2_^+^ -NTA agarose overnight. Solution with beads was placed in gravity column and washed with lysis solution, followed by 2 volumes of wash buffer (lysis buffer + 30 mM imidazole). Finally, protein was eluted with elution buffer (lysis buffer + 500 mM imidazole) and fractions were collected. Protein concentration was assessed by BCA assay and run on an SDS-page gel. The 3-4 highest concentration fractions were pooled and dialyzed against storage buffer (50 mM Tris-HCl pH=7.5, 150 mM NaCl, 1 mM DTT, 5% glycerol).

### Stable Isotope Labeling of Amino Acids in Cell Culture (SILAC)

Turnover experiments were performed using HEK293 cell lines stably expressing SIRT6 WT or centSIRT6. Cells were cultured for one week until confluent in MEM for SILAC (Thermo) supplemented with 10% dialyzed FBS (Gibco), L-glutamine, L-arginine ^13^C_6_ ^15^N_4_ (Cambridge Isotopes), and L-lysine ^13^C_6_ ^15^N_2_ (Cambridge Isotopes). The media was changed to normal culture medium and cell pellets were harvested at 0, 2, 4, 6, 8, 12, & 24 hours post-media change. Cell pellets were lysed in buffer containing 8 M urea, 75 mM NaCl, 50 mM Tris, pH 8.5, with protease inhibitor cocktail (Roche). Pellets were vortexed for 30s and sonicated 5x 10 s with 1 min rest on ice in between sonication steps. Lysate was centrifuged at 15,000 xg for 10 min and then supernatant was collected.

### SIRT6 deacylation activity

The substrate for this reaction is a synthetic TNF-derived peptide first developed by Schuster *et al* (Schuster, Roessler et al., 2016) was synthesized by Genscript. Reactions were performed with 150 mM NaCl, 20 mM Tris, 5% glycerol, 1mM β-mercaptoethanol, 2 µM SIRT6, and substrate concentration ranges of 7-1000 µM NAD^+^ and 2-88 µM peptide. Reactions were performed at 37°C and fluorescence was measured using 310/405 nm excitation/emission spectra on a Tecan Spark 20M plate reader. Readings were taken every 30s and initial rates were calculated from the relative fluorescence increase over a minimum of 6 min.

### Histone Analysis

Histones were purified from cumate SIRT6 fibroblasts (SIRT6 KO primary human foreskin fibroblasts constitutively expressing the catalytic subunit of telomerase as well as various alleles of SIRT6 via a cumate-inducible promoter) using an established acid extraction method followed by propionylation of lysines prior to trypsin digestion to enhance coverage (Shechter, Dormann et al., 2007, Sidoli et al., 2016). A data-independent analysis (DIA) mass spectrometry (MS) method was employed for quantitation of modified peptides after samples were separated by nano-LC using EpiProfile(Sidoli, Lin et al., 2015) or, alternatively, after direct infusion followed by a one-minute acquisition and analysis with EpiProfileLite (Sidoli, Kori et al., 2019).

### Analysis of SIRT6 and LMNA protein-protein interactions by mass spec

Cumate SIRT6 fibroblasts were used to compare interactomes for centSIRT6 and the wild type SIRT6. Nuclear extracts were prepared using hypotonic lysis buffer. Nuclei isolated from approximately 2.5x10^7^ cells were resuspended in one ml of extraction buffer (50 mM TrisHCl pH 7.5, 150 mM NaCl, 10% glycerol, 5 mM DTT, 0.25%NP-40, 1x Roche Complete protease inhibitors, as well as kinase/phosphatase inhibitors- 1 mM NaVO4, 10mM beta-glycerophosphate, 1mM sodium pyrophosphate, 1 mM NaF) and nuclei were disrupted by passage through a 27 gauge needle followed by sonication (3 pulses at constant 20% power) with a Branson Sonifier on ice. Next, samples were filtered through 0.2 μm MWCO filter to remove any insoluble material (and reduce non-specific binding). Proteins were quantified using the BCA assay and equal amount of wild type SIRT6 and centSIRT6 derived extract were divided into 5 replicates each. Three replicates of extract received 4 μg of anti-SIRT6 antibody (Cell Signaling #1248) or anti-LaminA (Lamin A/C-Millipore 05-714) and 75 μl of Miltenyi protein G μMACS magnetic particles. Two of the replicates were mixed with normal rabbit IgG (Cell Signaling #2729) as controls for non-specific binding to the particles. Samples were rotated at 4°C for 6 h followed by separation by the magnet. Samples were washed with 3 additional ml of extraction buffer and eluted with 100 μl of boiling elution buffer (5% SDS with 50 mM TrisHCl pH 7.5). After removal of SDS with S-columns (Protifi; Huntington, NY) and trypsin digestion, peptides were resuspended in MS-grade water and labeled with tandem mass tags (TMT 10-plex; Thermo Fisher). Samples were resolved by nano-electrospray ionization on an Orbitrap Fusion Lumos MS instrument (Thermo) in positive ion mode using a 30 cm home-made column packed with 1.8 μm C-18 beads to resolve the peptides. Solvent A was 0.1% formic acid and solvent B was 100% acetonitrile (CAN) with 0.1% formic acid. The length of the run was 2 h with a 90 min gradient. CID (35% collision energy) was used for MS2 fragmentation. HCD (60% collision energy) was used for MS3 detection of TMT groups. Other details of the run parameters may be found in the embedded run report of the RAW data file uploaded to the ProteomeXchange database. Peptide assignments were made using Proteome Discoverer and Sequest and MS3 ions were used for quantitation. False discovery rates (FDR) were estimated using a Decoy Database Search with Target FDR (strict) set to 0.01 and Target FDR (relaxed) set to 0.05. Validation was based on q-value. In the consensus step, ions with a co-isolation threshold above 30% were excluded. Normalization between replicates was achieved using the total protein approach (Wisniewski, 2017) where peptide counts for a single protein were divided by the sum of all proteins in the lane. Proteins appearing in specific antibodies (either anti-SIRT6 or anti-Lamin A) compared to normal IgG serum with Student *t*-test *p* <0.05) were considered as interactions. Similarly, proteins showing higher levels in centSIRT6 versus wild type SIRT6 were based upon *p*<0.05.

### SIRT6-NAD^+^ binding by Tryptophan Fluorescence

One μM of SIRT6 protein in 50 μl of buffer (50 mM TrisHCl pH=7.5 and 150 mM NaCl) was combined with a range of NAD^+^ (0-2 mM). Samples were then placed in a 384 Corning Flat Black plate and analyzed in a Tecan Spark 20M plate reader. Fluorescence was measured across 300-400 nm range in 2 nm steps using a 280 nm excitation. Background quenching by NAD^+^ of SIRT6 denatured by 7M urea was subtracted from the spectra as described previously(Pan, Muino et al., 2011).

### Deacetylase Assay

Three μg of SIRT6 protein was combined with 0.5 μg histones, 1 or 5 mM NAD^+^, 30 mM Tris-HCl pH=8, 4 mM MgCl_2_, 1mM DTT and ddH_2_O up to 50 μl. Designer nucleosomes with relevant PTMs were obtained from Epicypher. All reagents were prepared on ice and moved to 37°C for the duration of the incubation. Reaction was quenched with direct application of 1 volume of 2x Laemmli buffer with BME. Samples were boiled for 15 min before being run in Western analysis using anti-H3K9ac and anti-H3K18ac antibodies.

### PARP1 Ribosylation Assay

Five μg of SIRT6 protein was combined with 100 fm PARP1, 20 mM Tris-HCl pH=8, 10 μM ZnCl_2_, 10 μM MgCl_2_, 10% glycerol, 300 μM NAD^+^, 1 mM DTT, 0.1 μg/ml salmon sperm DNA and ddH_2_O up to 50 μl. All reagents were prepared on ice and moved to 30°C for 30 min. Reaction was quenched with direct application of 1 volume of 2x Laemmli buffer with BME. Samples were boiled for 15 min before being run in Western analysis using anti-PADPR antibody.

### SIRT6 Ribosylation Assay

Three μg of SIRT6 was combined with 50 mM Tris-HCl pH=7.5, 1 mM DTT, 10 μM ZnCl_2_, 150 mM NaCl, 25uM NAD^+^, 1 μl [^32^P]- NAD^+^ (800Ci/mmol, 5mCi/ml; Perkin Elmer BLU023x250UC) and ddH_2_O up to 20 μl. All reagents were combined on ice in a master mix except for SIRT6. Master mix was aliquoted into reaction tubes and SIRT6 was added and incubated at 37°C for 3h. Reaction were quenched with 1 volume of 2x Laemmli buffer with BME. Samples were boiled and then run on SDS-PAGE gel and then transferred to PDPF membrane. Self-ribosylation was assayed using phosphoimager.

### Thermostability

Four μM SIRT6 protein was combined with 1x SYPRO Orange dye in storage buffer (50 mM Tris-HCl pH=7.5, 150 mM NaCl, 1mM DTT, 5% glycerol) to 50 μl. Samples were placed in qRT-PCR plate and run on a BioRad CFX Connect Real Time machine using a Melt Curve protocol (30°C-75°C, 0.5°C steps in 10 s intervals).

### FRET

Mutagenesis to generate the centenarian mutant (N308K/A313S) was performed using NEB Q5 site-directed mutagenesis kit, following protocols for primer design.

HEK293-6E cells were transfected using 293fectin protocol (Thermo Fisher). The SIRT6 biosensors were expressed using mammalian expression vector pTT5 (National Research Council, Canada). After two days in culture, approximately 30 million were produced, generating transiently transfected cell lines expressing the biosensor at high levels. The cell line was maintained using F17 medium (Sigma) + (200nM/mL) GlutaMAX.

On each day of FRET assays, approximately 30 million cells from each transfection were harvested, washed three times in PBS with no Mg or Ca (Thermo Scientific; Waltham, MA) by centrifugation at 300 x g, filtered using 70-µm cell strainers (Corning; Corning, NY), and diluted to 1-2 million cells/mL using a Countess™ Automated cell counter (Invitrogen; Carlsbad, CA). On each day of experiments, cell viability was assessed using the trypan blue assay. After resuspension and dilution in PBS, the cells were continuously and gently stirred with a magnetic bar at room temperature to keep the cells in suspension and prevent clumping.

An in-depth description of the fluorescence instrumentation has been published previously(Schaaf, Peterson et al., 2017). For FLT measurements, the observed fluorescence waveform was convolved with the instrument response function to determine F(t) Eq.1, the average energy transfer efficiency E was calculated from the average FLTs of donor τ_*D*_ and donor-acceptor, τ_*DA*_, using Eq. 2, and the donor-acceptor distance R was calculated using Eq. 3.

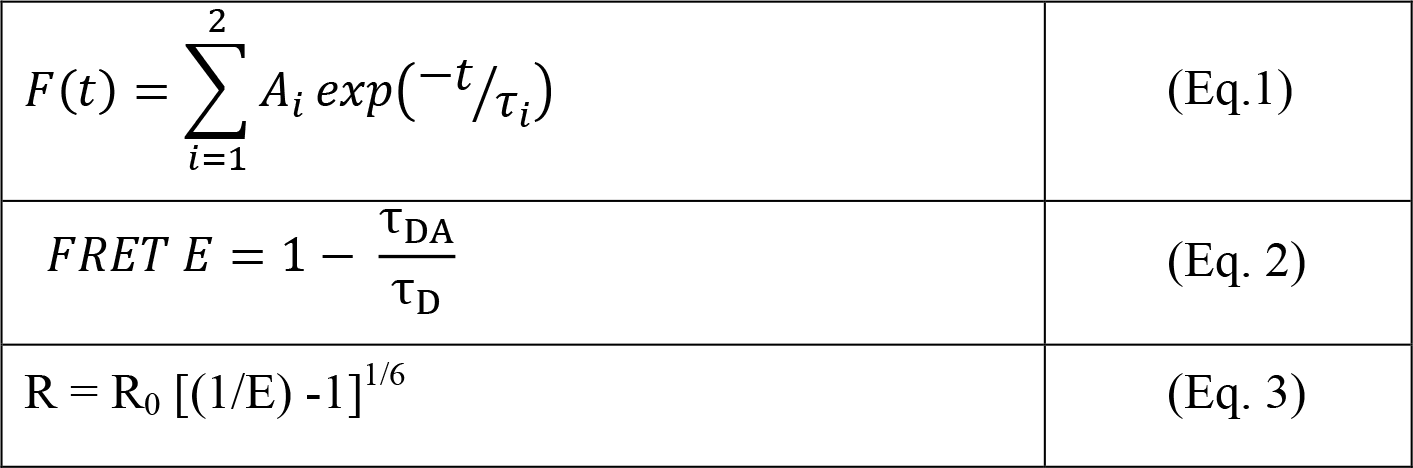

|  |  |
| --- | --- |
| $F(t) = \sum_{i=1}^2 A_i \exp(-t/\tau_i)$ | (Eq.1) |
| $FRET\ E = 1 - \frac{\tau_{DA}}{\tau_D}$ | (Eq. 2) |
| $R = R_0 [(1/E) - 1]^{1/6}$ | (Eq. 3) |

### Statistics

Unless otherwise noted, the Student’s *t*-test was used to determine statistical significance between groups. All tests were two-tailed and *p*-values were considered significant below a 0.05 threshold. Cell culture experiments utilized at least two separately derived cell lines for each genotype and were performed in triplicate unless noted otherwise. In vitro assays were done in triplicate using two or more separate protein isolation preps.

## DATA AVAILABILITY

### Lead Contact

Further information and requests for resources and reagents should be directed to and will be fulfilled by the lead contact, Vera Gorbunova

### Materials Availability

Plasmids and cell lines generated in this study are available upon request

### Data and Code Availability

All data needed to evaluate the conclusions in the paper are present in the paper and/or the Supplementary Materials. All sequencing data have been deposited in the Sequence Read Archive (SRA) under the bioproject number PRJNA669033, PRJNA669034 and PRJNA669037. MS data files will be uploaded to PRIDE/ProteomeXchange data repository and the Digital Object Identifier number will be provided prior to publication.

## Supplementary Data

Data S1: Mass spectrometry data for histone PTM quantification Data S2: Mass spectrometry data for SIRT6 interacting proteins Data S3: Mass spectrometry data for LMNA interacting proteins

## ACKNOWLEDGEMENTS

We would like to extend a special thanks to Dr. Ivan Matic for providing the mADPr antibodies. We are thankful to Kevin Welle, Jennifer Hryhorenko of Mass Spectrometry Resource Lab for their help and advice. Funding was provided by the US National Institute of Health.

## AUTHOR CONTRIBUTIONS

Conceptualization: YS, VG, AS, MS, Methodology: MS, EE, VG, AS, YS

Investigation: MS, JY, JG, TMS, LZ, MG, SLY, AP, MVM, JH, SR, AT, YZ, AH, GA, NB, AW, KM, BAG, DDT, PDR, SE, MZ, GT, MG, EE

Visualization: MS, VG, AS, YS

Funding acquisition: VG, AS, YS, PDR, BAG Project administration: VG, AS, YS Supervision: VG, AS, YS

Writing – original draft: MS

Writing – review & editing: MS, JY, YS, JV, VG, AS, EE

## EXPANDED VIEW FIGURES

**Figure EV1.**
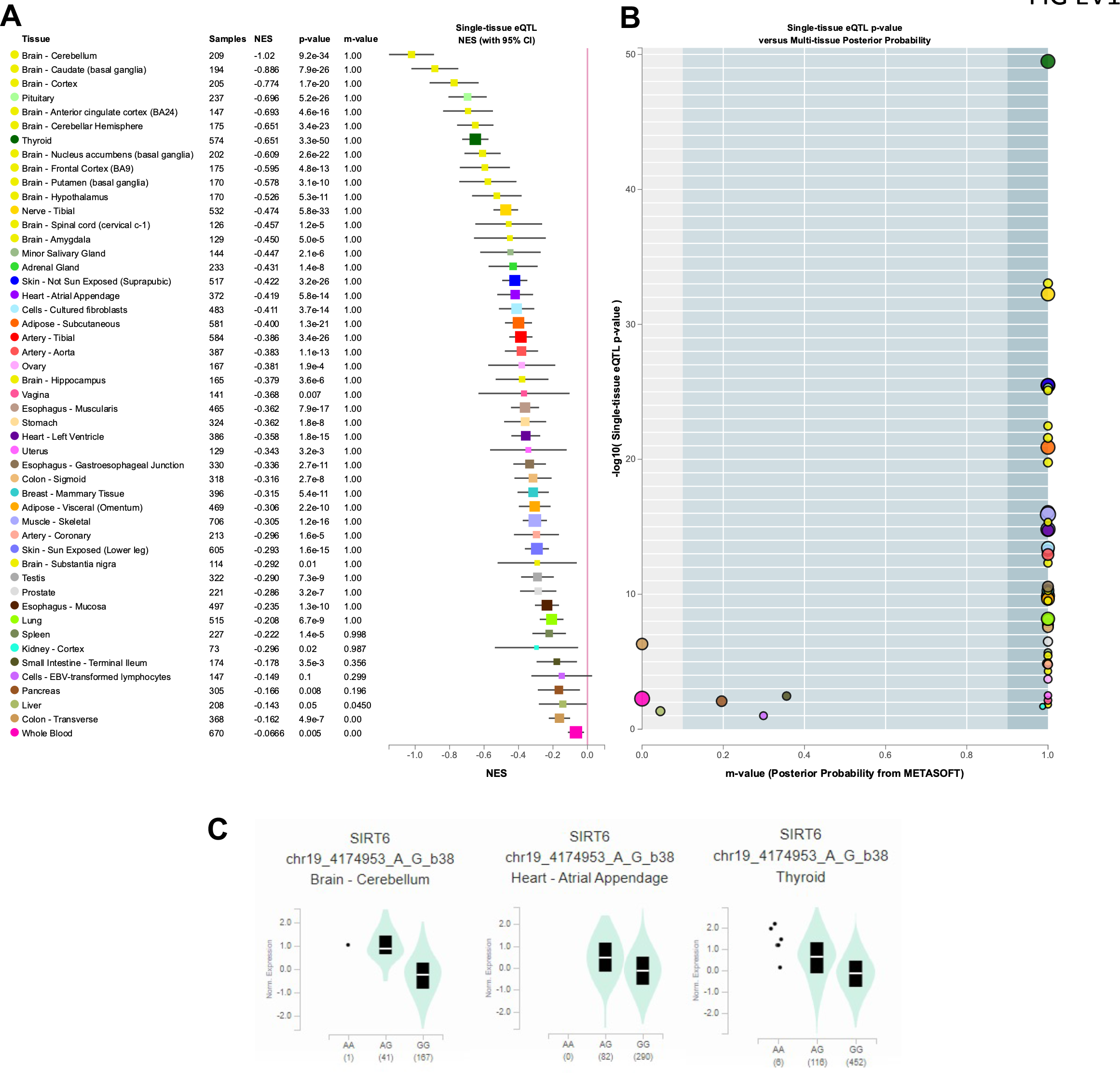
rs350845 (chr19:4174953:A:G) is a *cis*-eQTL for *SIRT6* upregulation across multiple tissues. **A** GTEx Tissue types are shown sorted by m-value which is the posterior probability that an eQTL exists for *SIRT6* in a specific tissue from a cross-tissue meta-analysis. NES = Normalized Effect Size based on single-tissue analysis; p-value is from a t-test comparing observed NES to a null within a single-tissue. **B** Single tissue eQTL p-value vs. m-values. **C** Violin plots showing *SIRT6* expression differences across three example tissues with m-value = 1. Reference allele is G and alternate allele is A, and each panel shows the normalized expression of *SIRT6* in a different tissue. Values beneath genotypes represent number of individuals.

**Figure EV2:**
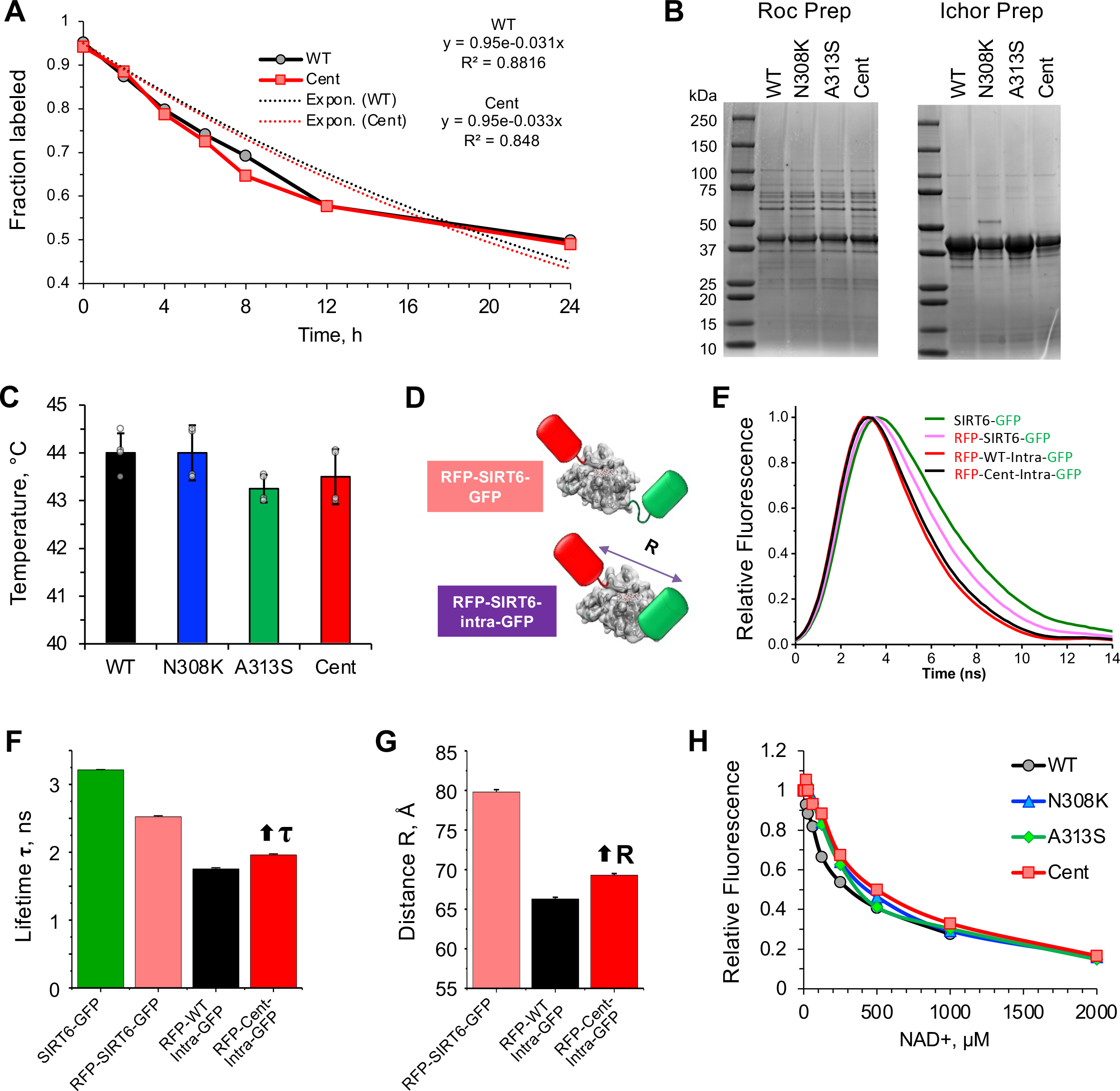
Turnover rate and biochemical properties of purified SIRT6 proteins. **A** Protein turn-over rate. SILAC analysis on HEK293 cells expressing SIRT6 variants. **B** Coomassie staining of SIRT6 purified from Rosetta-Gami *E. coli* cells. Proteins were purified at the University of Rochester (Roc) as well as the Ichor Therapeutics facility in Ithaca. All *in vitro* experiments were performed with both protein preparations and the data was consistent between the two preps. **C** Thermostability of purified SIRT6 proteins. Data represents two replicates with two technical replicates each using SIRT6 from the Roc and Ichor preps. **D** FRET biosensor design. Top: RFP-SIRT6-GFP with RFP (acceptor) and GFP (donor) fused to N- and C-termini of SIRT6. Bottom: RFP-SIRT6-intra-GFP with GFP inserted internally within SIRT6 after D206. **E** Representative Fluorescence lifetime (FLT) signals acquired from four different transiently transfected cell lines. **F** Fluorescence lifetime τ measured from FLT signals by fitting the data to Eq. 1 (see Methods). Each of the two different SIRT6 biosensors shows a decreased fluorescence lifetime compared to the GFP-only control, indicating highly significant FRET. Compared with the biosensor with C-terminal GFP (pink), the intra-GFP biosensor (black) shows a greater lifetime change (more FRET, shorter distance R, more structural sensitivity). The centSIRT6 of this biosensor was also tested (red). It shows a significant increase in lifetime τ (1.96 ± 0.02 ns), compared to the wild type SIRT6 (1.75 ± 0.01 ns). Error bars indicate SEM (n = 3-5). **G** The FRET efficiency E and the distance R were determined from lifetime data (Eqs. 2 and 3, Methods), revealing that the distance R, measure for the intra-GFP biosensor, was significantly greater for the centSIRT6 by 3.0 ± 0.4 Å. **H** Tryptophan fluorescence curves for SIRT6 variants with titrated concentrations of NAD^+^.

**Figure EV3:**
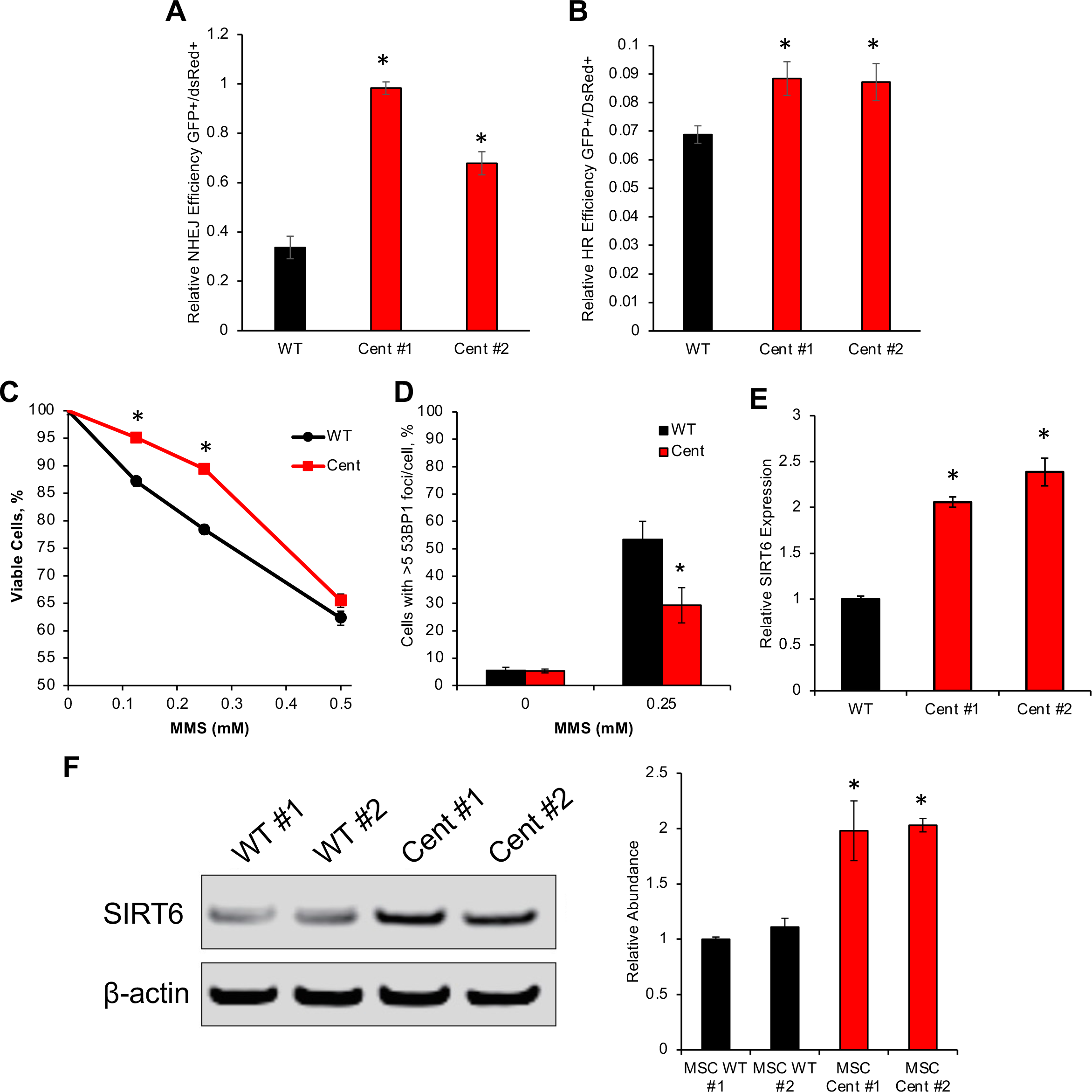
HeLa histone deacetylation by recombinant SIRT6, and SIRT6 expression in cumate-inducible cell lines. **A** *In vitro* deacetylation rates of purified SIRT6 proteins with histones purified from HeLa cells. **B** SIRT6 expression in cumate-inducible telomerase-immortalized HCA2 human fibroblast cell lines. SIRT6 alleles were integrated in the genome of SIRT6 knockout HCA2 cells using PiggyBac Transposon Vector System. Different dosages of cumate and resulting SIRT6 abundance were used to determine the dose needed to achieve equivalent SIRT6 expression for each cell line. Cells were normalized by count and total protein. Subsequent experiments utilizing cumate-inducible SIRT6 fibroblasts were controlled for SIRT6 abundance using these data. For each cell line, the red box indicates the corresponding cumate dosage for equivalent expression (WT=60 μg/ml, N308K=30 μg/ml, A313S=30 μg/ml, and Cent=7.5 μg/ml). These concentrations were used in experiments with these cells. **C** qRT-PCR expression analysis of SIRT6 using standardized dosages of cumate (dosage corresponds to red boxes in Fig EV3B **D** Coomassie stained SDS-PAGE of histone preparations for quantitative mass spectrometry

**Figure EV4:**
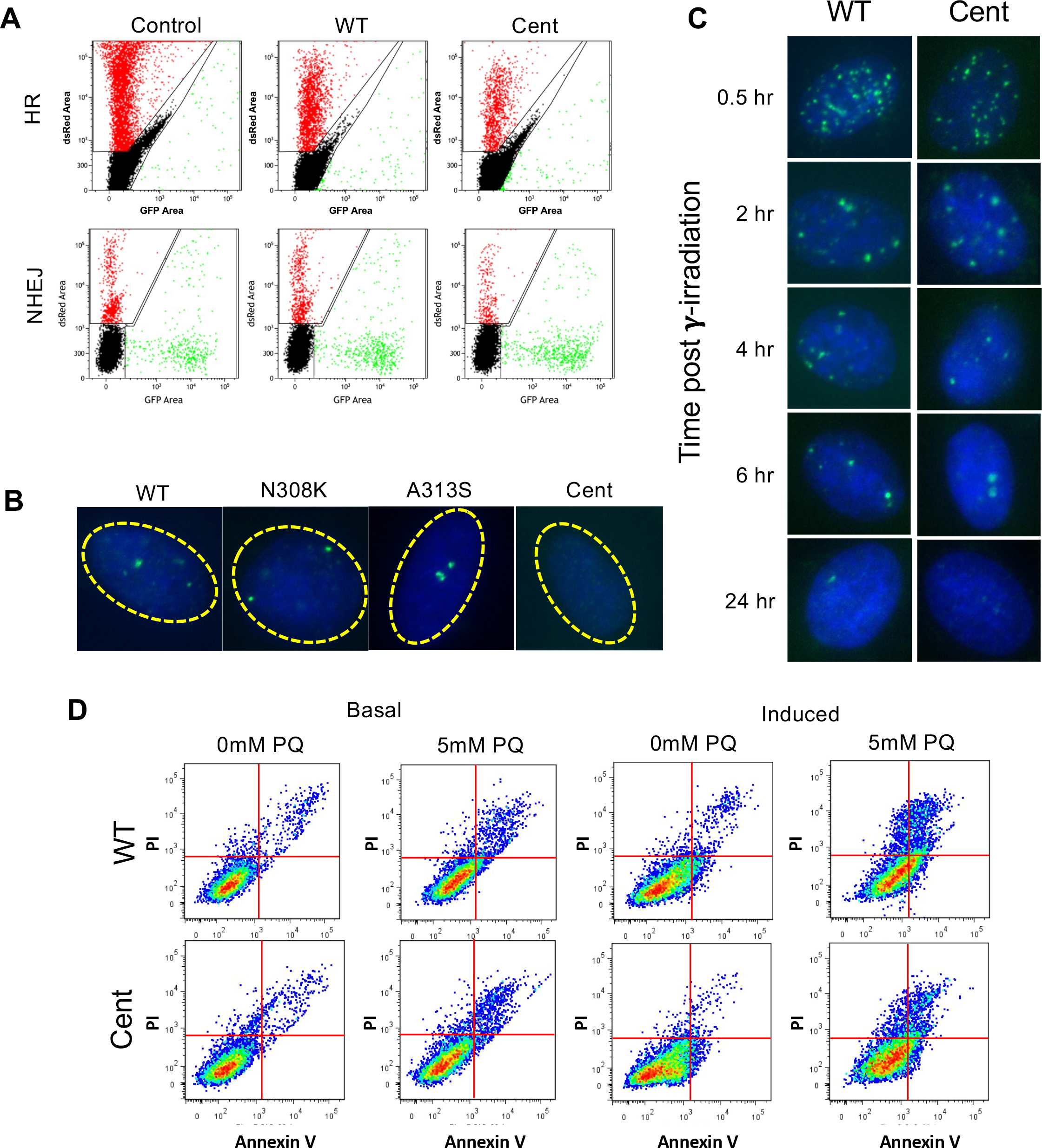
Quantification of SIRT6 expression and FACS gating for DNA repair assays. **A** Representative images of FACS analysis for DNA DSB repair assays (Fig 4B, C). **B** Representative images of basal γH2AX immunostaining (Fig 4D). **C** Representative images of γH2AX immunostaining after γ-irradiation (Fig 4E). **D** Representative images of FACS traces for paraquat sensitivity assay (Fig 4F, G).

**Figure EV5:**
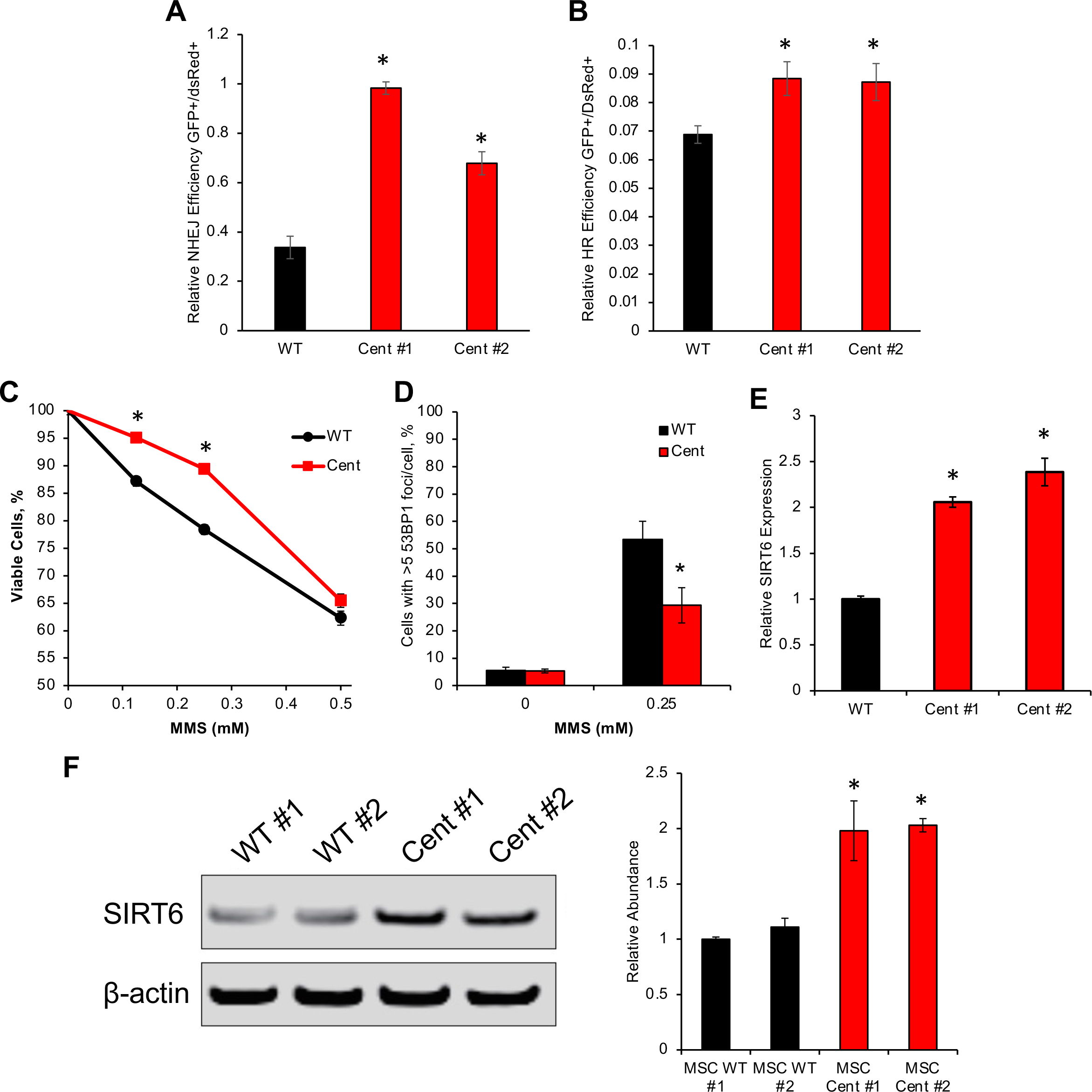
Analysis of CRISPR human MSC cell lines. **A, B** DNA double strand repair efficiency in wild type and centSIRT6 hMSCs. DSB repair reporter constructs were integrated into hMSCs. After 72 hr recovery, reactivation of the GFP reporter was measured by flow cytometry. Stimulation of NHEJ or HR was calculated as radio of GFP^+^/DsRed^+^ positive cells. **C** Cell viability in MMS-treated wild type and centSIRT6 hMSCs. hMSCs were treated with MMS for 48 hours and cell viability was evaluated by MTS assay. Data were normalized to the control group (0 mM). n = 6. **D** Immunofluorescence staining of 53BP1 in wild type and centSIRT6 hMSCs under MMS treatment. Numbers of 53BP1 foci in the nuclei of wild type and centSIRT6 hMSCs with or without MMS (0.25 mM) treatment were quantified. >600 nuclei from 10 images were scored. **E** qRT-PCR analysis of SIRT6 expression in wild type and centSIRT6 hMSCs (P2). Data normalized to Actin and were presented as mean ± SEM, NS, not significant. **F** Western blot analysis of SIRT6 in wild type and centSIRT6 hMSCs. β-Actin was used as a loading control and used to normalize quantification. Error bars represent s.d. Significance was determined by Student’s *t*-test, two tailed. Asterisk indicate *p*<0.05

